# Adaptive evolution of odorant receptors is associated with elaborations of social organization in ants

**DOI:** 10.1101/2025.08.20.670513

**Authors:** Yoann Pellen, Joel Vizueta, Erin Beck, Juergen Leibig, Lukas Schrader, Eyal Privman

## Abstract

Cooperation in social insect colonies depends on complex chemical communication, requiring a large array of chemosensory receptors. Ant odorant receptors (ORs) were dramatically expanded compared to other insects, most notably in the “9-exon” subfamily, which was implicated in responding to cuticular hydrocarbons, a major class of signalling compounds. These observations indicate adaptive evolution of olfactory functions, but this process was never studied in the context of the evolution of specific sociobiological traits. The Global Ant Genomics Alliance has compiled 163 high-quality ant genomes, enabling detailed study of OR evolution in unprecedented detail. Analysing 55,068 ORs across the phylogeny, we tested for association between sociobiological traits and adaptive evolution of ORs, including gene duplication and adaptive sequence evolution. We identified strong enrichment of positive selection on 9-exon ORs in the ancestor of the formicoid clade, which evolved larger colonies and greater reproductive division of labour. This result indicates a key role of chemical communication in the early evolution of complex social organization. We also observed enrichment of positive selection on 9-exon ORs associated with the recent evolution of continuous worker polymorphism in multiple lineages. Surprisingly, the evolution of other sociobiological traits was associated with reduced positive selection on ORs. These results suggest that worker polymorphism involves more extensive adaptation of chemical communication compared to other aspects of ant sociobiology. By analysing the most comprehensive OR dataset to date, we provide new insights into the specific context in which ORs played a major role in the elaboration of social traits in ants.

## Introduction

Animal species vary in their sociality and the complexity of social systems. Communication systems allows social animals to reliably assess the behaviour of others and to respond to it adaptively (reviewed in Peckre et al. 2019). Freeberg *et al*. (2012) discussed variation in complexity of sociality and communication systems and defined complex societies as those in which individuals frequently interact in many different contexts with many different individuals. They defined complex communicative systems as those that contain a large number of structurally and functionally distinct elements (i.e., a large repertoire of signals) or possess high information content. The evolution of greater social complexity was found to be associated with complexity of vocal communication in various mammals, including subterranean rodents (Vanden Hole et al. 2014) and mongooses (Manser et al. 2014). In social insects, chemical communication is a vital function mediating many aspects of colony organisation. Colonies of social insects such as ants are characterised by elaborate cooperation among nestmates. Communication is central to the efficient coordination of necessary activities, including foraging for food, caring for the queen and brood, nest construction and defence. Cooperation is critically dependent on recognition of colony members, known as nestmate recognition, which is mediated by chemical cues such as cuticular hydrocarbons (CHCs). CHCs are a diverse group of long-chain hydrocarbons thought to be a major component of the chemical cues used for recognition, including sexual signals, nestmate recognition, and regulation of reproductive division of labor, which require classifying individuals based on their CHC profile (van Zweden and d’Ettorre 2010; Van Oystaeyen et al. 2014; Blomquist et al. 2020; Liebig and Amsalem 2025). Extensive research showed that CHC profiles are more complex in ants relative to other insects and revealed rapid evolution that generated great diversity among ant species (Martin and Drijfhout 2009), and also to some extent in the broader Aculeata (ants, bees, and wasps) (Kather and Martin 2015).

The perception of chemical signals by ants relies on a large repertoire of odorant receptors (ORs), as well as other smaller chemosensory receptor families (Bargmann 2006; Bear et al. 2016; Robertson 2019). ORs are critical for recognition and communication in ants (Trible et al. 2017; Yan et al. 2017). In particular, the 9-exon (“9e”) subfamily of ORs was suggested to harbour many of the receptors that respond specifically to the diverse array of CHCs displayed by ants (Engsontia et al. 2015; Zhou et al. 2015; McKenzie et al. 2016; Pask et al. 2017; Slone et al. 2017). This clade of the OR gene tree is dramatically expanded in ants, accounting for about a third of all ORs (C.D. Smith et al. 2011; C.R. Smith et al. 2011; Zhou et al. 2012; Zhou et al. 2015; McKenzie et al. 2016). Pask et al. (2017) characterized 22 ORs from the 9e clade and showed that they are narrowly tuned to respond to specific CHCs. Most of the 9e ORs display worker-biased expression, in line with a role in social coordination in the colony (Zhou et al. 2012). These ORs are also mainly expressed in the ventral half of the antennal club that comes in contact with other ants when they antennate each other for nestmate recognition, in line with these being receptors of non-volatile contact pheromones such as CHCs (McKenzie et al. 2016). In social insects, discrimination against non-colony members is especially important to avoid any form of colony exploitation. Therefore, strong selection on these discrimination abilities is expected.

The transition from solitary to social organisation in ants was accompanied by a dramatic expansion of the OR gene family (C.D. Smith et al. 2011; Zhou et al. 2012). Generally, ORs evolve by rapid gene duplications and losses, in a so-called “birth-and-death” process of gene family evolution (Ohno 1970; Demuth and Hahn 2009; Innan and Kondrashov 2010; Mendivil Ramos and Ferrier 2012). Newly generated paralogs may undergo positive selection for novel adaptive olfactory functions (Smadja et al. 2009). Positive selection on ant OR sequences is especially focused on the putative ligand binding site, in line with tuning for new ligands in different species (Saad et al. 2018). Therefore, the expansion of ORs in ants indicates adaptive evolution, which may have contributed to the evolution of sociality. This may be limited to the ants, as recent work found no evidence of a general relationship between the evolution of eusociality in Hymenoptera and the expansion of the OR repertoire (Gautam et al. 2024). However, another study did find evidence of episodic positive diversifying selection in expanded ORs subfamilies, mainly in the 9e clade, in *Polistes* paper wasps (Legan et al. 2021). Ant genomes harbour hundreds of ORs, and up to 503 ORs reported for the clonal raider ant *Ooceraea biroi* (McKenzie and Kronauer 2018). Studies have also shown that the reductive social behaviour in social parasite species (e.g. *Formica selysi, Acromyrmex ameliae, Lasius cf. spathepus, Tetramorium atratulum*) was linked to a loss of OR genes (Schrader et al. 2021; Jongepier et al. 2022; Vizueta et al. 2025). Many ant ORs show evidence of positive selection on their protein sequence (Engsontia et al. 2015; Zhou et al. 2015; Saad et al. 2018) making them attractive candidate genes for the study of adaptive evolution of chemical communication in ants. In total, Saad et al. (2018) reported positive selection in 7.2% of the tested OR gene tree branches (252 out of 3507). Recent species-specific OR gene duplications were followed by a higher rate of positive selection compared to ORs that were not recently duplicated (Engsontia et al. 2015; Saad et al. 2018), as expected for adaptive evolution in the birth-and-death model of gene family evolution.

The study of ant ORs greatly benefits from the recent high-quality genomic dataset produced by the Global Ant Genomics Alliance (GAGA; Vizueta et al., 2025). GAGA sequenced genomes of 145 species of ants, which, in addition to 18 published high-quality genomes, represent 27 tribes in 12 subfamilies. Most of these genomes were produced using long-read sequencing that resulted in highly contiguous genome assemblies. This is especially important for accurate and complete assembly and annotation of the large tandem arrays of OR genes. Previous studies based on short-read genome assemblies revealed only a partial picture, typically underestimating the number of ORs. For example, the first genome assembly of *Camponotus floridanus* that was based on Illumina short-read sequencing allowed the annotation of 352 complete OR genes and 55 partial genes (Zhou et al. 2012), while the more recent PacBio long-read assembly (Shields et al. 2018) resulted in annotations of 466 complete ORs and 61 partial genes. For this species, we also generated long-read RNA sequencing data (using the Iso-Seq protocol) to validate the predicted transcript sequences and evaluate the accuracy of our gene annotations. The genomic dataset produced by GAGA allows us to study the dynamic evolution of the OR gene family in ants with greater resolution and precision than was previously possible.

The genome-wide analysis of single-copy gene families in the GAGA genomes (Vizueta et al. 2025) revealed a high rate of positive selection on protein sequences in the branch leading to the ancestor of formicoid ants, a clade containing most extant ant species, which split from the poneroid ants ca. 125-145 Mya. The formicoid clade exhibits the most elaborated forms of reproductive division of labour, large colony size (Dornhaus et al. 2012), worker polymorphism (Oster and Wilson 1978), and extended queen longevity (Keller and Genoud 1997). This enrichment of positive selection in this branch is consistent with previous work published by Romiguier et al. (2022) which identified this branch as a hotspot for genes under positive selection. The GAGA analyses also revealed expansions in multi-gene families in the ancestor of ants. Many of these expanded gene families were implicated in chemoperception (22 families) and biosynthesis of CHCs (17 families). These results are in line with the hypothesis that the evolution of sociality in the ant ancestor involved the evolution of sophisticated chemical communication.

Here, we focus on the evolutionary dynamics of ant ORs and explore whether positive selection on ORs is linked to the evolution of traits such as diet, colony size, worker polymorphism, and worker reproduction. We hypothesized that the evolution of such traits involved the evolution more complex chemical communication systems, in terms of a larger number of distinct signals (larger vocabulary) and/or the evolution of relatively simple signals into more complex, multi-component signals (i.e. signalling greater information content; e.g. a more complex CHC profile). We predicted that the perception of more complex signalling require a larger number of ORs. Thus, ant OR are expected to have undergone adaptive evolution involving expansion via gene duplications and positive selection, which generated a larger array of receptors with distinct ligand specificities. We tested our hypotheses using sets of representative species across the entire phylogeny, focusing on branches with inferred trait shifts. We identified enrichment of positive selection on the 9e ORs subfamily in the formicoid ancestor. Surprisingly, more recent branches with a transition to bigger colony sizes were depleted for positive selection, and that depletion was mostly in the non 9e OR subfamilies. We generally observed more depletion than enrichment in the recent branches we tested. However, positive selection on 9e ORs was enriched in branches of lineages that evolved continuous worker polymorphism. Our analyses leverage the large-scale data from the GAGA project to reveal a complex pattern of adaptive evolution of ant ORs associated with some, but not all, elaborations of sociobiology in the ant phylogeny.

## Materials and methods

### Traits of interest

The GAGA project included the compilation of a trait table with information on social and ecological traits for each species from literature and experts, including for example their habitat, diet, foraging stratum, colony size, queen number, worker polymorphism, etc. We focused on traits that we hypothesised to be related to ORs evolution: log10 maximum colony size (here after referred to only as colony size), diet, worker polymorphism, and worker reproductive functions. Unfortunately, we could not include queen number in our analysis because of uncertainty in the ancestral reconstruction of this trait. These traits require sensory function such as recognition and discrimination, whether of nestmates or of food sources, as well as complex social interactions that require complex signalling. Therefore, ORs can be expected to play a major role in the evolution and elaboration of these traits. Our analysis was designed to test for association between adaptive evolution of ORs and the evolution of greater social complexity such as larger colonies or worker polymorphism. Ancestral state reconstruction for each trait across the ant phylogeny was conducted using the R package *ape* (version 5.7-1; Paradis and Schliep, 2019) evaluating the best fitting model by using the Akaike information criterion (AIC). Further details can be found in Vizueta et al. (2025). Trait reconstructions are plotted on the phylogeny in Supplementary Figures S1 to S4. We used these reconstructions to identify branches with inferred trait changes, which we refer to as “transition branches” (marked on Figure 2). We hypothesized that there was positive selection on ORs related to the evolution of the transitioning traits in these branches. For example, we identified branches with a transition from an omnivorous to a herbivorous diet (e.g. in the ancestor of *Messor* harvester ants) and branches where colony size increased (e.g. in the ancestor of *Lasius* and *Myrmecocystus*).

The number of species analysed here coupled with the large number of ORs annotated in each of them resulted in a dataset of 55,068 OR genes. Tests for positive selection using dN/dS models are not feasible for such a large gene tree. Therefore, we focused the positive selection analyses on nine clades with small subsets of species (four to nine species in each; 75 species in total). The species in each clade were selected as representative of the phylogeny that allow us to test for positive selection on the transition branches, i.e., at least two species were selected that define each of the nodes above and below the branch. We also tested the three most ancestral branches of the formicoid clade (rectangles in Figure 2) using a selection of ten representative species across the entire ant phylogeny. The first of these branches is the ancestor of all formicoid ants, the second is the branch after the split of the *Dorylinae* subfamily, and the third is the ancestor of *Formicinae* and *Myrmicinae* after the split of the *Myrmeciinae, Pseudomyrmicinae, Aneuretinae* and *Dolichoderinae* subfamilies.

### Genome sequences

Most of the species in the GAGA dataset have been sequenced using PacBio long-read sequencing (125 out of 163 species), which is advantageous relative to Illumina short-read sequencing for the assembly and annotation of repetitive regions such as the typically large arrays of tandem-duplicated OR genes. Twenty species, including some rare species like *Leptanilla*, had limited sample biomass and could only be sequenced with stLFR (single-tube long fragment read), a sequencing technique that requires fewer DNA amount than PacBio. Scaffolding of contigs into whole chromosomes was achieved for 12 species using HiC, a technique that captures genome-wide chromatin interactions (Belton et al., 2012). The published genomes of 18 additional species were retrieved from NCBI, giving a total of 163 genomes. Among the 75 genomes used in our analyses, 65 were sequenced by PacBio and seven sequenced by stLFR. Three of the non-GAGA genomes were based on 454 sequencing, which is an obsolete short-read technology that is expected to result in more partial and missing ORs. Further details about the genome sequencing and assembly can be found in Vizueta et al. (2025).

### Antennal transcriptome Iso-Seq

For the purpose of total RNA isolation, founding queens of *Camponotus floridanus* ants were collected from the Florida Keys in 2018 and raised to mature colonies in the laboratory at 25 °C. A total of twelve biological samples were prepared from six colonies, consisting of two caste groups – major and minor workers (one sample of each from each colony). To obtain sufficient material, each of the twelve samples was a pool of sixty antennae harvested from thirty individual ants belonging to the corresponding major or minor caste group. Total RNA was extracted from the collected antennae using the standard Trizol/Chloroform precipitation method. The purified RNA was dissolved in RNase-free water and stored at -80°C until shipped on dry ice to the California Institute for Quantitative Biosciences (QB3) Vincent J. Coates Genomics Sequencing Laboratory for isoform sequencing (Iso-Seq) using the Pacbio Kinnex protocol, in one SMRTcell of a Revio sequencer. The sequencing resulted in a total of 10,878,248 reads, 97.49% of which had full arrays, and a mean array size of 7.94 (Supplementary figure S5).

### Gene annotation

OR gene annotation was done using the *HAPpy-ABCENTH* pipeline (https://github.com/biorover/HAPpy-ABCENTH). The *ABCENTH* (Annotation Based on Conserved Exons Noticed Through Homology) program is an algorithm for building gene models that was designed for multigene families like ORs, with high sequence divergence but a conserved exon structure. *ABCENTH* builds HMM profiles separately for each exon based on known protein coding genes, which increases sensitivity and precision. *ABCENTH* then looks for splice sites of the candidate exons according to the expected exon length and phase.

The genes used as queries were taken from manually curated ant ORs of the species: *Harpegnathos saltator, Stigmatomma sp, Ooceraea biroi, Camponotus floridanus* and *Solenopsis invicta*. Subsequently, novel OR annotations were evaluated by comparing them to OR sequences from *Drosophila melanogaster* and the five abovementioned ants. We identified the OR co-receptor (ORCO) in each species. Annotated ORs were classified in three categories: complete genes with the expected exon structure, pseudogenes with in-frame stop codons, and partial or fragmented annotations with incomplete exon structures, which can be caused by pseudogenization, fragmented assembly or assembly and annotation errors.

The accuracy of the annotation pipeline was tested through a leave-one-out approach, by re-running it on the genome of *Camponotus floridanus*, which was previously annotated and has and antennal RNA-sequencing data available (data from Zhou *et al*. (2012)). We reannotated this genome using ORs from *Ooceraea biroi, Harpegnathos saltator* and *Solenopsis invicta* as queries. The old annotations by Zhou *et al*. were generated on the *C. floridanus* genome version 3.5 (based on Illumina short-read sequencing), while we used here version 7.5 (based on PacBio long-read sequencing), which is expected to result in differences. The old annotations had 352 ORs that were considered complete (average length of 393 amino acids) although 122 of them were missing the start codon. Without manual intervention, the new annotation found 451 complete ORs (average length of 392 amino acids). A blast of the old sequences on the new show an average identity of 98.12%, and 0.31% of gaps, with 351 of the 352 old sequences having a complete hit (more than 95% of its length) in the new annotation. Furthermore, out of the 122 sequences missing a start codon in the Zhou sequences, 92 now have it in the new annotation. We then compared our gene models to the Iso-Seq data we generated from Pacbio HiFi sequencing of antennal RNA from *C. floridanus* workers. Blasting our new sequences on the Iso-Seq data, we found that 449 of our 451 complete genes had a nearly perfect hit (more than 99% coverage and more than 99% identity) and are supported on average by 12,432 reads (average identity of 99.76%, and 0.07% of gaps), while fragmented genes are supported by 1,298 reads and ORs flagged as pseudogenes by 3,218 reads.

### Phylogenetic analysis

The large number of sequences – 55,068 complete ORs – made it impossible to analyse the full dataset jointly. We started by assigning all ORs in all species into 29 subfamilies, as defined in McKenzie et *al*. (2016). Each subfamily of ORs contains all orthologs from across the ant phylogeny. Therefore, for each of the clades of species we chose, each of the 29 OR subfamilies was analysed separately, and the results were pooled together for statistical analysis (see subsection “Testing for association” bellow). ORs were assigned to a subfamily using *BLASTP* (version 2.11.0; Altschul et al., 1990), based on the top hit to sequences classified in the published subfamilies. 98% of the ORs (54,051 out of 55,068) had their top five hits within the same subfamily. The 9e subfamily comprises 19,528 sequences in our dataset, so it was further subdivided into six monophyletic clades that had 100% bootstrap support (we also observed issues in the alignment of these clades to each other, which would result in false positives in the dN/dS tests).

We constructed a multiple sequence alignment and tree for each OR subfamily and subset of species chosen for the positive selection analysis. Nucleotide coding sequences were used to build codon alignments using *Guidance2* (version 2.02; Sela et al. 2015) with the aligner *PRANK* (version 140110; Löytynoja and Goldman, 2005). Unreliably aligned codons were masked at a cutoff on the *Guidance* confidence scores that was chosen ad hoc for each alignment. We started by masking each alignment at a threshold of 0.8, and if more than 20% of the codons were masked we lowered the threshold by 0.1 and tested the percentage of masked codons again. We repeated the operation until we achieved a maximum of 20% masked codons, to avoid losing too much information for the inference of positive selection. This procedure was demonstrated as good practice for controlling the false positive rate of the branch-site test for positive selection (Privman et al. 2012). The OR gene trees were built using *RAxML* (version 8.1.15; Stamatakis, 2014) with the translated sequence alignments as input, using the PROTCALG model, the LG rate matrix, and 100 bootstraps repeats.

### Positive selection inference

Positive selection was inferred using the branch-site test for positive selection (Zhang 2005) as implemented in *Godon* (Davydov et al. 2019), using the M0 model to estimate branch length, four categories for codon rate variation, and testing all branches. A likelihood ratio test was conducted for each gene tree branch contrasting between the null model, not allowing positive selection, and the alternative model, allowing for positive selection in that branch. The likelihood ratio test p-values were then corrected for multiple testing using the Benjamini-Hochberg (1995) method.

### Testing for association between trait evolution and OR sequence evolution

We developed a novel method for mapping positive selection results from genes trees of multi-copy gene families to the species tree, which is described in detail in our methodological paper (Pellen and Privman 2025). Briefly, we tested for enrichment of positive selection on ORs in species tree branches where we inferred a change in our traits of interest (the “transition” branches). The *Generax* (version 1.1.0; Morel et al., 2020) algorithm for reconciliation between gene trees and species trees was used to map branches of the gene trees onto branches of the species tree, with the UndatedDL reconciliation model. This step allows mapping of gene tree branches that were tested for positive selection to the species tree branch that we wish to test for enrichment. We then used a hypergeometric test to test for enrichment of positive selection (i.e. gene tree branches having *Godon* p-value < 0.05) on the species tree “transition” branches. Since we expect it to be more likely that positive selection occurred in a long branch than in a short branch, we normalized the number of gene tree branches by a factor defined as their branch length relative to the average gene tree branch length of the studied clade. This normalized number of branches was used as the size of the population that is tested for enrichment in the hypergeometric test. The ratio of observed/expected frequency of gene tree branches with positive selection were calculated to determine by what factor was the transition branch enriched or depleted for positive selection.

We pooled the positive selection results before conducting the enrichment test. The positive selection tests were performed on separate datasets for each of the 29 OR subfamily and for each of the 9 clade of species, that is 261 datasets in total. Therefore, we pooled all of the results for a given type of trait change (e.g. larger colony size) across all datasets before testing for enrichment of positive selection (Figure 1).

**Figure 1:**
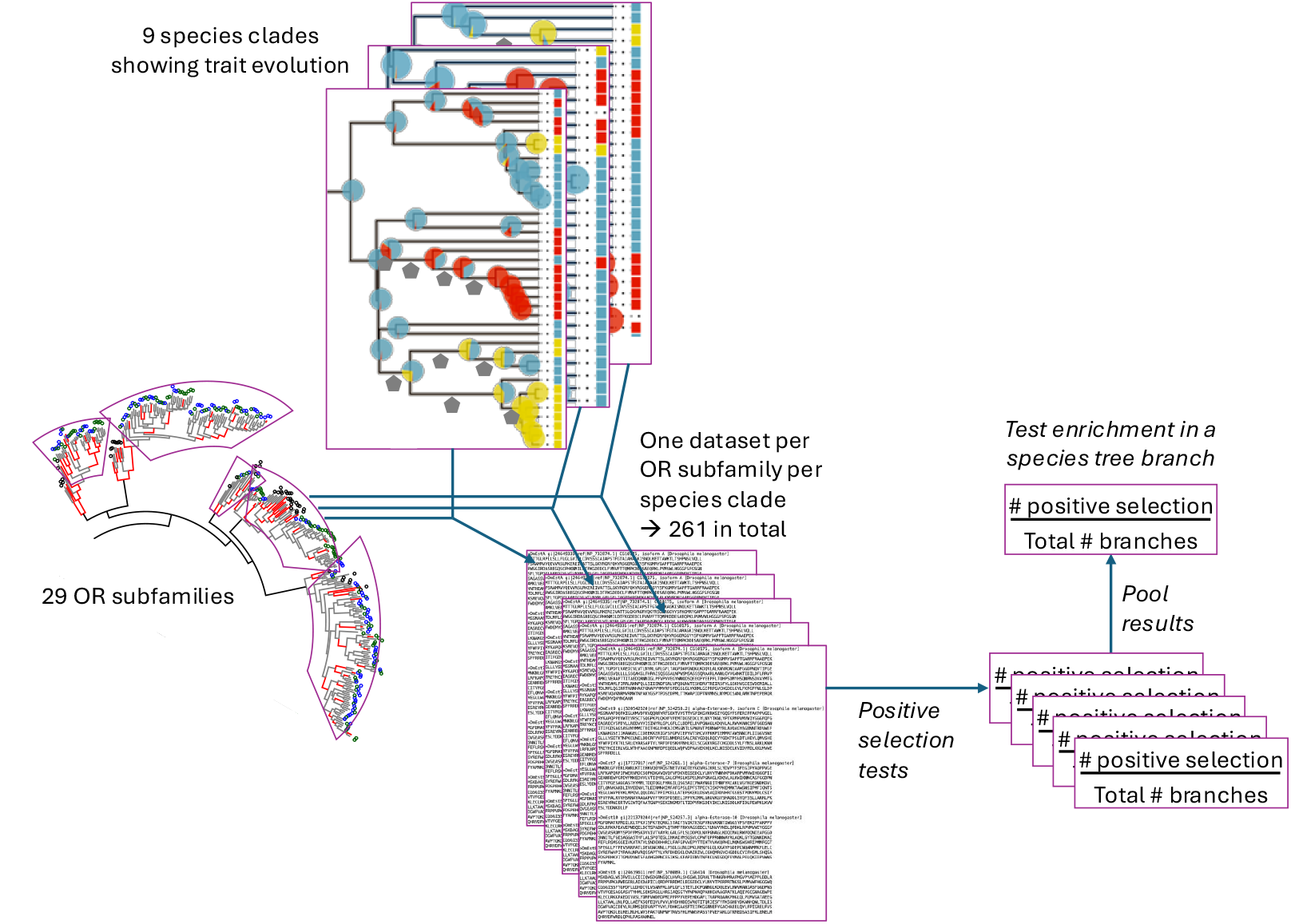
Analysis scheme for subdividing and pooling the data before the statistical analysis.

### OR gene family expansions and contractions

To analyse the evolution of OR genes numbers, we applied the BDI stochastic model implemented in the *BadiRate* program (Librado et al. 2012). To avoid bias due to genome incompleteness or fragmentation, we excluded the 20 low quality genomes (short-read based with N50 < 500Kb), except for the *Leptanilla* outgroup, and added *Apis mellifera* as an outgroup. We then applied a global rates model (BDI-GR-CSP in *BadiRate*; Sankoff parsimony) to the numbers of complete OR genes. We note that the number of ORs reconstructed for the ant ancestor may be an underestimate due to *Leptanilla sp*., the most basal ant species with only 151 complete ORs annotated. This genome has a lower quality compared to the rest of the dataset, as it is sequenced with the stLFR method only. It has an N50 size of 0.13 Mb, while the average for the dataset is 4.55 Mb. This low assembly contiguity may result in many ORs appearing to be partial genes which are not counted in the final dataset.

We also conducted a Phylogenetic Generalized Least Squares (PGLS) analysis using the *pgls* function in the R package *caper* (version 1.0.3, https://CRAN.R-project.org/package=caper) using the dated ant phylogeny. Again, we excluded the 20 species with lower quality genome assemblies by using *drop*.*tip* in the R package *treeio* to prune the species tree. We used PGLS to fit regression models between traits and ORs numbers (Supplementary Table S2). OR subfamilies with mostly single copy genes were removed from the final results to reduce spurious correlations.

## Results

### The odorant receptor repertoire of ants

The total number of ORs found through the annotation of 163 ant genomes was 66,169. This count includes partial ORs, which can be missing some terminal and/or internal parts of the full-length sequence. The percentage of partial ORs in each species varied from 4% to 56% (Supplementary Table S1) and was related to the contiguity of the genome assembly (scaffold N50 size), with fragmented assemblies having more partial ORs (Pearson correlation *r* = -0.28 between N50 scaffold size and the percentage of partial ORs, *p*-value = 0.00074). For example, *Gigantiops destructor* has the lowest N50 (8,895 bp) and the highest proportion of partial ORs (56%). Low quality genomes (N50 < 500Kb) were therefore excluded from analyses regarding gene numbers as they are expected to have biased under-estimates of OR numbers. We excluded partial ORs from further analyses, which left a total of 55,068 complete ORs across all 163 genomes, and 50,657 ORs without the low-quality genomes, ranging from 90 (*Tetramorium “Anergates” atratulus*) to 687 (*Pseudoneoponera rufipes*), with an average of 338 ORs per species (Figure 2). The ancestral reconstruction of OR gene numbers, using *Apis mellifera* as an outgroup, estimated 378 ORs for the ancestor of ants not including *Leptanilla*, which was excluded due to its low-quality genome (see inferred ancestral numbers in Supplementary Figure S6). The number of ORs expanded to 455 in the ancestor of the formicoid clade, while there was no further significant expansion in the poneroids ancestor (with 375 ORs).

**Figure 2:**
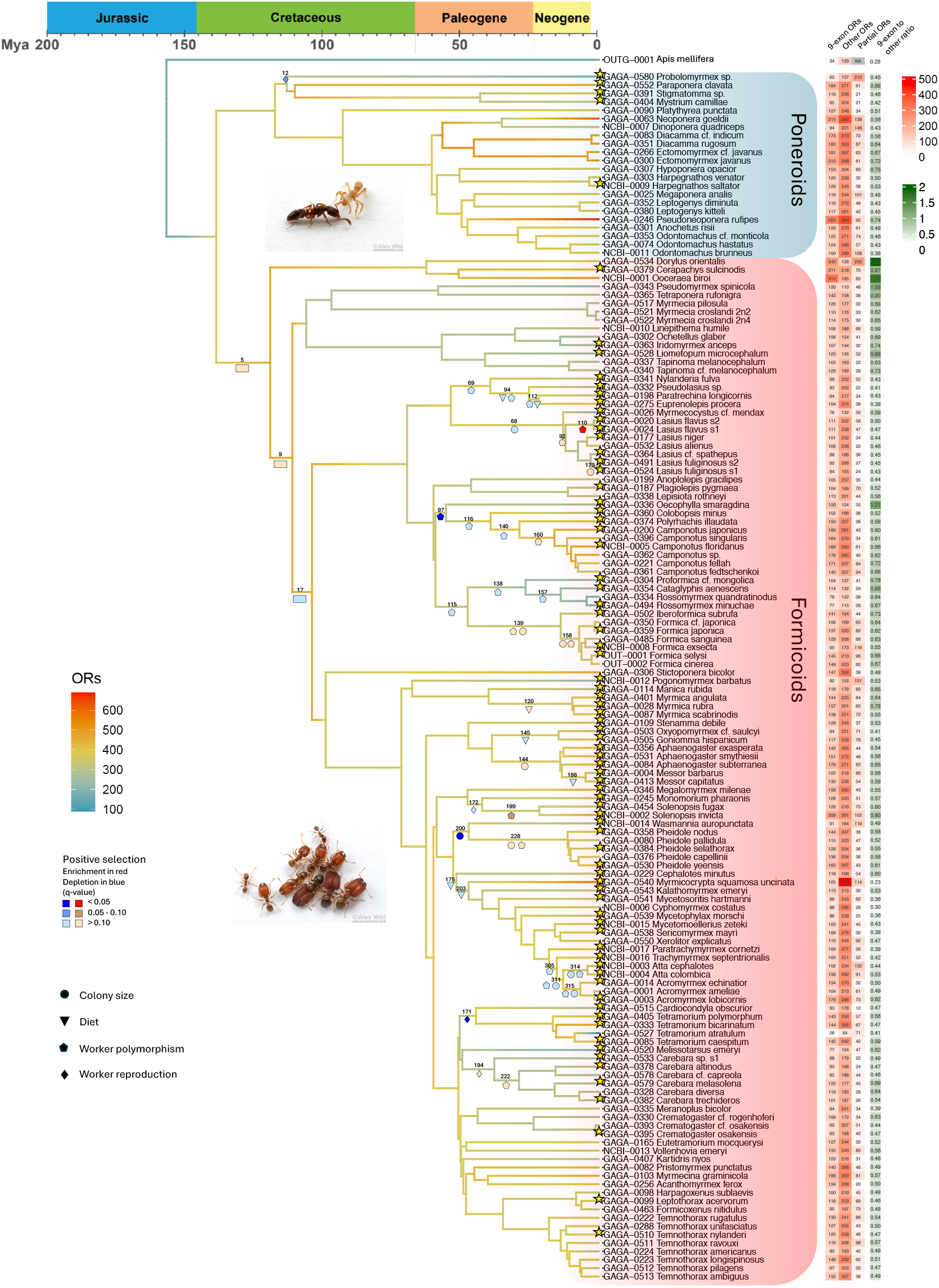
OR evolution along the ant phylogeny, including ancestral reconstruction of OR numbers and positive selection enrichment tests. Branch colour indicates the inferred ancestral number of ORs (see inferred ancestral numbers in Supplementary Figure S6). Coloured shapes mark branches with trait shifts that were tested for enrichment of positive selection on ORs. Shape colour indicates the statistical significance of enrichment or depletion based on the multiple testing corrected *p*-values (*q*-values). The species used in the positive selection analysis are marked with stars. The heatmap shows the number of complete ORs for each species partitioned to the 9-exon clade and the rest of the ORs, as well as partial genes.

**Figure 3:**
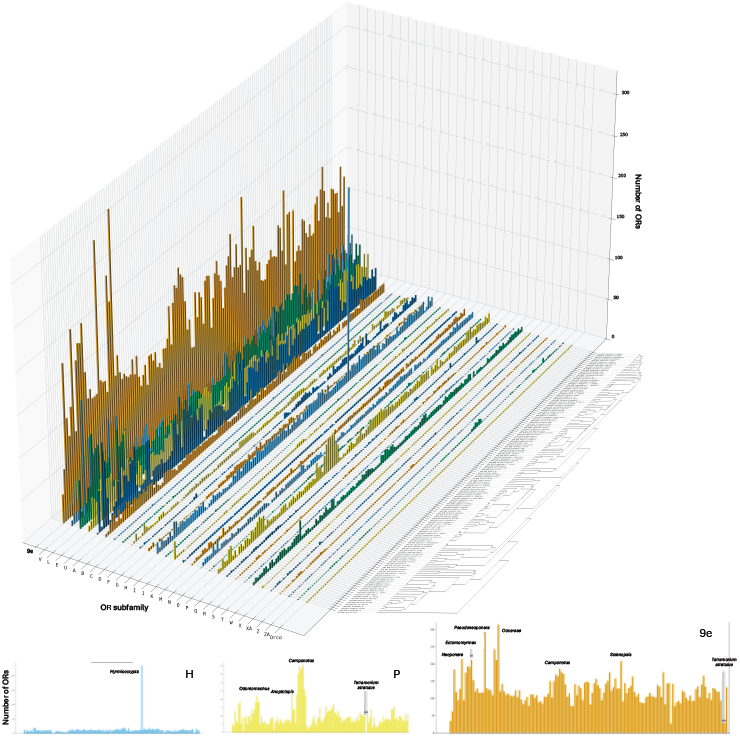
Subfamily composition of the OR superfamily across the ant phylogeny. The plotted data include only complete ORs, and exclude the species with low-quality genomes.

We observed substantial variation in OR numbers along the phylogeny. Most species have fewer 9e ORs than non 9e ORs (Figure 2). The only exceptions being the small clade of army ants. *Ooceraea biroi* is the most extreme case with 315 9e ORs against 185 non 9e, as previously reported for this species (McKenzie et al. 2016; McKenzie and Kronauer 2018). Conversely, we note the *Myrmicocrypta* species as an extreme outlier for the number of non 9e ORs, with 458. Gene numbers of many OR subfamilies vary dramatically across ant species, most notably in the 9e subfamily and the other four largest subfamilies (E, L, U, V). Looking at the variation in the 9e ORs (Supplementary Figure S7), one species stands out, the workerless social parasite *Tetramorium atratulus*, which has the smallest number of 9e ORs. This species also shows drastic reductions in the other large OR subfamilies and is the only ant species that completely lost the medium-sized subfamilies P, T, M, and N. In subfamily P (Supplementary Figure S8), we noted a clade-specific expansion in the clade that includes *Camponotus, Polyrachis* and *Anoplolepis* (with smaller number in *Oecophylla*), and ponerine clade-specific expansions in *Neoponera* and in the clade of *Odontomachus* and *Leptogenys*. In subfamily H (Supplementary Figure S9), *Myrmicocrypta* stands out with 236 ORs, against an average of 12 over the rest of the ants. This seems to be due to extensive tandem duplications of subfamily H ORs that are restricted to this species (Supplementary Figure S10).

### Correlation of OR gene numbers with social and ecological traits

The elaboration of sociobiological traits may require an extended OR repertoire to enable more sophisticated chemical communication in ant colonies. Therefore, we predicted a correlation between the number of ORs and traits such as colony size or worker polymorphism. We also expected that adaptation for different diets may be associated with the OR gene numbers. Therefore, we tested for an association between the total number of ORs or the size of OR subfamilies and specific traits of interest using a PGLS analysis: colony size, social parasitism, worker polymorphism and worker reproductive functions, and diet (Figure 4A and Supplementary Table S2). We observed an expansion of the 9e OR subfamily in species with larger colony size (*q*-value = 0.012), while another large subfamily, V, showed a significant contraction with bigger colony size (*q*-value = 0.012). Our study confirms previously described significant contractions of the OR gene family associated with the evolution of social parasitism (Schrader et al. 2021; Jongepier et al. 2022; Vizueta et al. 2025). This contraction was in the total number of ORs (*q*-value = 0.000015) and specifically in the 9e (*q*-value = 0.000027) as well as several other OR subfamilies. The only exception is subfamily N that was expanded in social parasites (*q*-value = 0.070). We further focused the analysis on inquilines, the more extreme form of social parasitic species that live in the nest of another ant species. Inquilines show an even stronger contraction (*q*-value = 0.000000027), which is particularly striking in *Tetramorium atratulum*, a workerless inquiline species which retained only 90 complete ORs, 26 of which are in the 9e subfamily compared to an average of 120 9e ORs in other ant species (Figure 4 and Supplementary Table S1). Contrary to our expectations, we found no significant correlation between OR gene numbers and worker polymorphism, worker reproduction, or diet.

**Figure 4:**
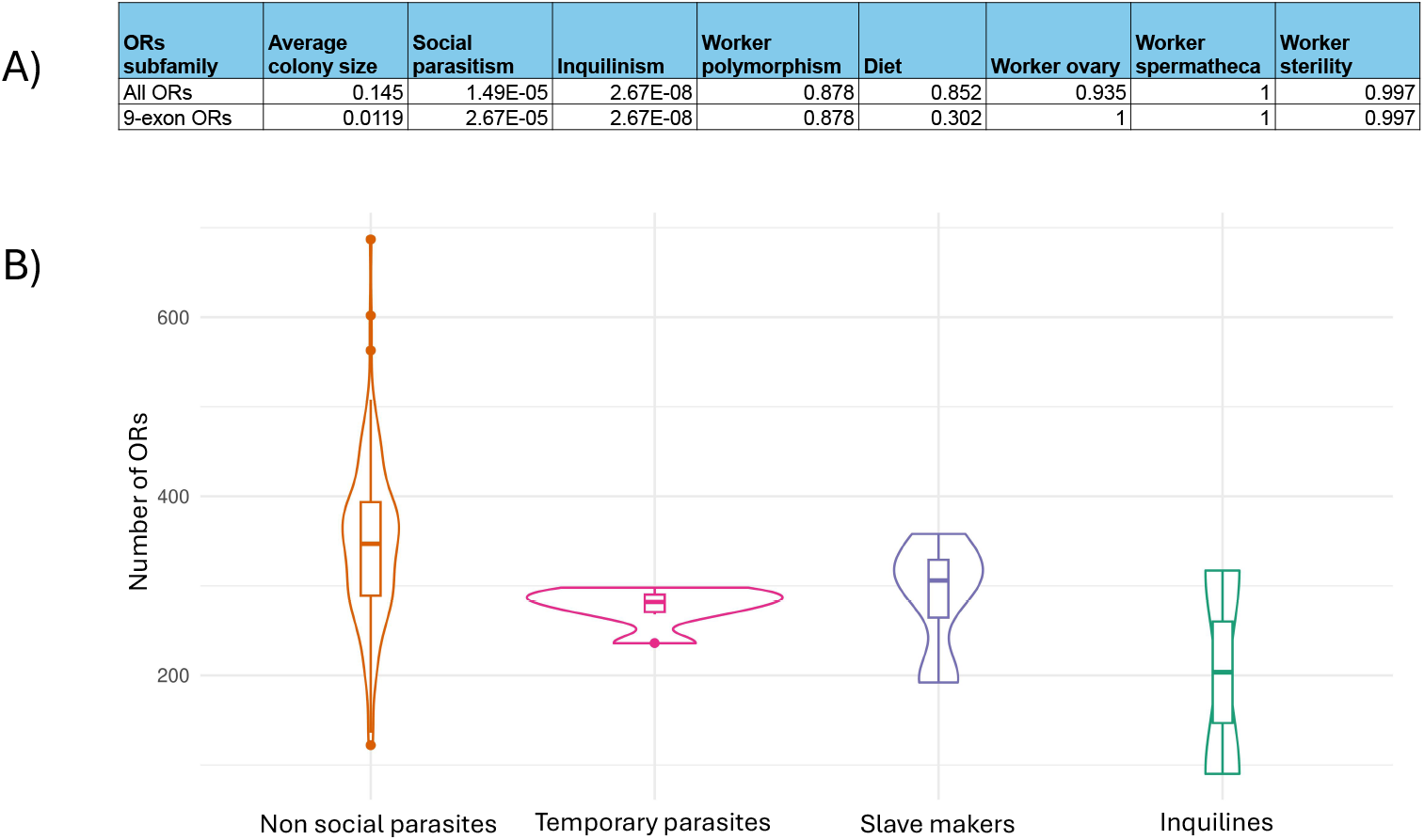
A) Phylogenetic Generalized Least Squares (PGLS) tests for association of gene family expansion or contraction with chosen traits, for the whole OR superfamily and for the 9-exon subfamily. Results are presented after correction for multiple testing (*q*-values). B) Number of ORs for species that are either not socially parasitic, temporary parasites, slave makers or inquilines.

### Positive selection on ORs

We used the branch-site test for positive selection to infer adaptive evolution of OR protein sequences in individual branches of the OR gene tree. We then evaluated whether species tree branches were enriched or depleted in positive selection on ORs (see Methods). We focused this analysis on subsets of the species phylogeny that allowed testing branches where we observed shifts in our chosen target traits (“transition branches”) based on ancestral trait reconstruction: colony size, worker reproduction, worker polymorphism, and diet (marked by shapes in Figure 2). This analysis involved testing branches of the gene trees of each OR subfamily for each subset of species, 249 gene trees in total. Out of a total of 89,802 gene tree branches tested across all our focus clades, positive selection was detected in 652 branches (0.7%), with a false discovery rate of 10% after correction for multiple testing.

To obtain better statistical representation for testing the association of positive selection with the evolution of a certain trait, we pooled the results for transition branches showing the same trait change. For example, we pooled the ten different species tree branches that showed a convergent increase in colony size to conduct a single test for enrichment of positive selection; and we pooled eight species tree branches where continuous worker polymorphism evolved for a single enrichment test. We generally observed depletion in positive selection associated with most trait transitions (Figure 5). Despite being statistically significant, the depletion seen for increased colony size has a negligible effect size (0.998-fold depletion, *p*-value=0.036). The evolution of continuous worker polymorphism was the exception, with a significant enrichment of positive selection (1.18-fold enrichment, *p*-value=0.007). Also the ancestral branches of the formicoid clade were enriched for positive selection (1.075-fold enrichment, *p*-value=0.0476).

**Figure 5:**
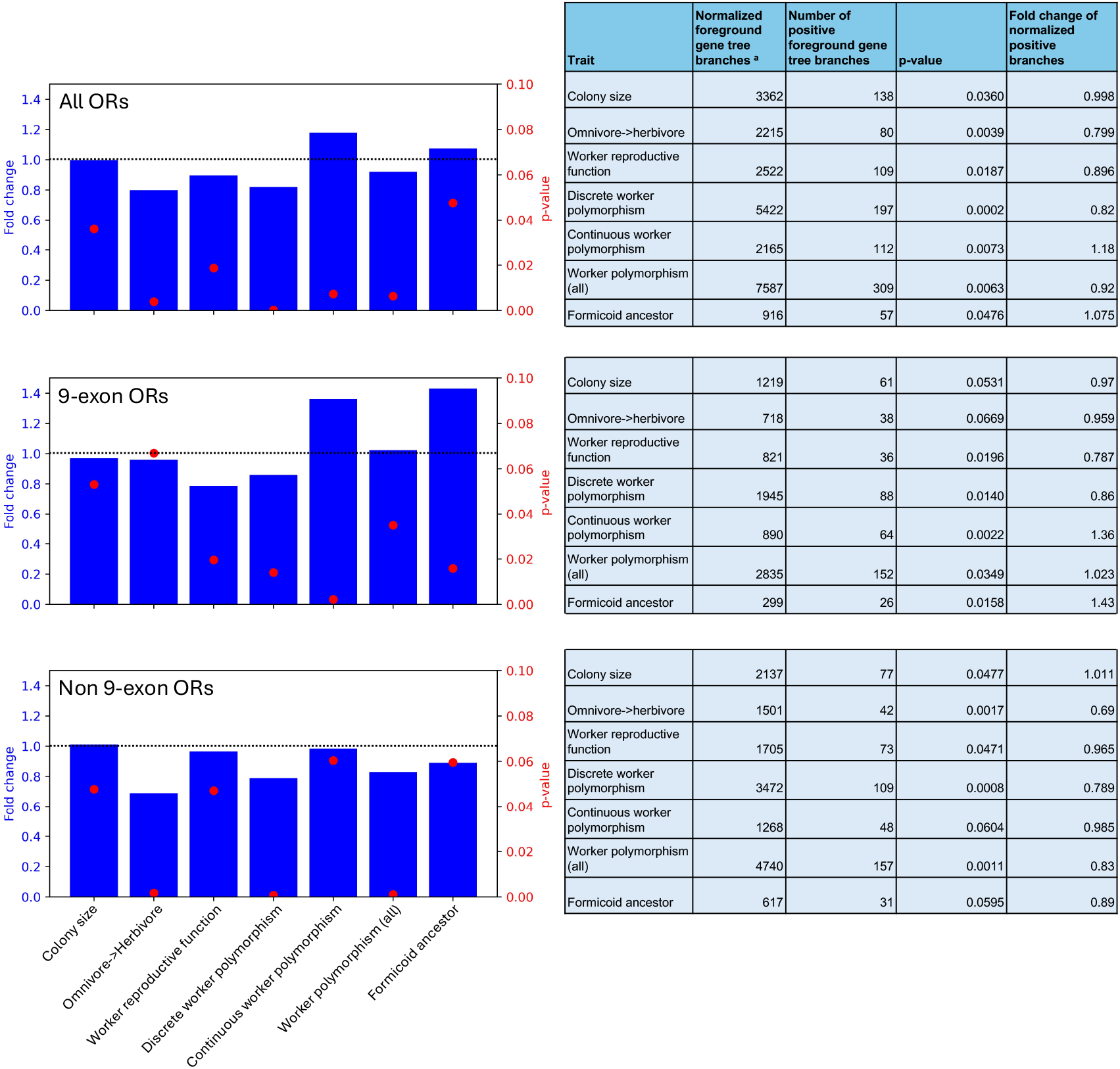
Enrichment for positive selection on ORs associated with trait evolution. Results of the hypergeometric test for enrichment of branches under positive selection (considering Godon’s p-values under 0.05 as positive) for the pooled branches exhibiting the same change in trait. a: Length-normalized number of gene tree branches mapped on the species tree branch that is tested.

We were especially interested in positive selection in the 9e subfamily because it was suggested to be involved in social communication. We found that the enrichment in the formicoid ancestor was driven by the 9e subfamily (1.43-fold enrichment, *p*-value=0.016; Figure 5), while non 9e ORs were depleted for positive selection. A similar pattern was observed for the species tree branches associated with the evolution of continuous worker polymorphism (1.36-fold enrichment in the 9e ORs, *p*-value=0.0022). In contrast, the shift to discrete polymorphism showed an opposite effect, with a significant depletion (0.863-fold depletion, *p*-value=0.014). In general, the enrichment or depletion in the 9e ORs mostly follows the pattern for total ORs but shows a stronger effect. The exception to this being the depletion associated with changes in diet, from omnivory to herbivory, which was mainly associated with the non 9e ORs.

In addition to the pooled results, we also inspected individual transition branches. Across all ORs subfamilies, we found significant enrichment of positive selection in 2 branches and depletion in 4 branches, out of 37 transition branches in the species that were tested (*q*-value < 0.10; Supplementary Table S3). For the three ancestral formicoid branches, individual branches were marginally significant (p-value 0.12). The two more ancient branches were enriched and the most recent one was depleted for positive selection. Some branches display especially strong depletion or enrichment for positive selection. For example, the common ancestor of *Wasmannia* and *Pheidole*, where increased colony size was inferred (Supplementary Figure S1), was strongly depleted for positive selection (0.534-fold depletion, *q*-value=0.03262). Out of the 14 branches in which discrete polymorphism evolved (Supplementary Figure S3), four were enriched and ten depleted for positive selection. Among these, the common ancestor of the clade including *Oecophylla* and *Camponotus* was strongly depleted (0.477-fold depletion, *q*-value=0.03262) of positive selection. The common ancestor of *Cardiocondyla* and *Tetramorium*, showing a loss of functional ovaries in workers (Supplementary Figure S4), was strongly depleted for positive selection (0.596-fold depletion, *q*-value=0.02461).

## Discussion

The high quality of the GAGA genomes and wide coverage across the ant phylogeny, coupled with an annotation pipeline specifically tailored for ant ORs, resulted in 50,657 ORs across 143 species, a more comprehensive and complete dataset for the study of the evolution of ant ORs than any previously generated dataset. We used this novel gene set to infer ancestral OR repertoire sizes, reconstruct gene trees for all ant OR subfamilies, and test for adaptive evolution of ORs that may be associated with important sociobiological and ecological traits of ants.

Our tests found only 0.7% of branches under positive selection, which is low compared to previous studies that tested for positive selection in ant ORs (Zhou et al. 2015; Saad et al. 2018). This may be because we focused mainly on relatively recent events and short branches in the ant phylogeny, whereas previous studies analysed only a few species representing different ant subfamilies, making the tested branches much longer. It could also be because we tested many more branches, forcing us to be more stringent in our FDR correction.

Our results confirmed previous analyses, finding evidence of an ancestral expansion of the OR gene family in ants (C.D. Smith et al. 2011; Engsontia et al. 2015; Mier et al. 2022), and especially the 9e subfamily. We also observed further expansion in the formicoid ancestor, from 378 to 455 ORs, consistent with the further elaboration of sociobiological traits inferred in this lineage. Genome-wide analyses also inferred significant expansions in the common ancestor of ants in gene families implicated in chemical communication, including chemoperception and synthesis of cuticular hydrocarbons (Vizueta et al. 2025). These ancestral expansions suggest that the communication system of ants evolved greater complexity, analogous to the evolution of complex vocalization in mammals (Manser et al. 2014; Vanden Hole et al. 2014). Molecular evolutionary analyses of ORs cannot distinguish between the evolution of a larger number of distinct signals (each may be detected by a single OR) and the evolution of signals with greater complexity, such as multi-component CHC profiles (may require a combinatorial coding involving multiple ORs). The expansion of OR gene numbers is in line with either explanation or a combination of both.

Previous studies of adaptive evolution in ant genomes identified an enrichment in positive selection on the branch of the ancestor of all formicoid ants (Romiguier et al. 2022; Vizueta et al. 2025), but these analyses were limited to single-copy genes, and did not include large gene families such as ORs. This is because the study of adaptive evolution in multi-copy gene families requires complex reconciliation of gene trees with the species tree, to account for gene duplications and losses (Page and Charleston 1998). Here we put in first evidence for enrichment of positive selection on the OR gene family in the formicoid ancestor. This pattern is driven by the 9e subfamily of ORs that is highly enriched for positive selection, while non 9e ORs are actually depleted. This is in-line with the hypothesis that 9e ORs are responsible for social communication functions in ants, including their putative role in nestmate recognition, which is supported by their specificity for cuticular hydrocarbons and their expression in workers (Engsontia et al. 2015; Zhou et al. 2015; McKenzie et al. 2016; Pask et al. 2017; Slone et al. 2017; Couto et al. 2023; Marty et al. 2025). We propose that as the ancestor of formicoid ants evolved various elaborate sociobiological traits such as larger colonies and greater specialization of the non-reproductive castes, they evolved additional complexity in their chemical communication, including new 9e ORs that respond to chemical signals like CHCs shaped by positive selection. The rise in communicative complexity could translate in both the development of multimodal signals where a single information can be transmitted through different forms (Hölldobler 1999; Renyard and Gries 2025), but also in more uses for a single type of signal, like pheromones used for alarms, trails marking or signalling infections and diseases (Jackson and Ratnieks 2006; Dawson et al. 2025).

Overall, we observed more depletions in positive selection than enrichments in the transition branches we tested, especially in the most recent ones. We observed depletion of positive selection associated with increased colony size, complementing the findings by Vizueta et al. (2025) of genome-wide relaxation of purifying selection in species with increased colony size. Colony size was suggested to be a major driver in the evolution of ants, with many other social traits depending on it, including worker polymorphism (Bell-Roberts et al. 2024; Vizueta et al. 2025). The genome-wide analysis by Vizueta et al. revealed intensified selection associated with the recent loss of worker ovaries in multiple formicoid lineages, but predominantly relaxation of selection associated with the evolution of worker polymorphism. Our analysis found depletion of positive selection on ORs associated with the loss of worker reproductive functions. While we found that evolution of discrete polymorphism shows a depletion in positive selection, we detected enrichment of positive selection associated with the evolution of continuous polymorphism. This intriguing pattern would be interesting to investigate further, as it could imply that the evolution of different forms of worker polymorphism involved different mechanisms of chemical communication. The differences between our results and the analyses by Vizueta et al. may be attributed to the very different nature of the genes analysed: while we studied the fast-evolving multi-gene OR family, their analyses were limited to conserved single-copy gene families. The significant cases of enrichment of positive selection were most strongly represented in the 9e subfamily. It is interesting to note that transition to herbivory shows the opposite pattern with the depletion more pronounced in the non 9e ORs, a result that is in line with the hypothesis that non-social olfactory functions are mediated by other OR subfamilies. We interpret this result as reduced rate of adaptive evolution in foraging olfactory functions, which may be explained by the need for perception of more diverse odours in the ancestral state of omnivory. Conversely, there was no significant association between diet and the number of ORs. This surprising result suggests that species keep a similar number of ORs for foraging for different food sources, even if the rate of adaptive evolution in these genes is different. Overall, our results indicate that only the evolution of certain worker traits involved a detectable signature of positive selection on ORs, whereas the elaboration of other sociobiological traits was associated with less adaptive evolution of ORs. When considering OR evolution as a birth-and-death process including gene family expansions and contractions and positive selection on OR sequences, detailed analyses may reveal differentiated patterns, involving adaptive evolution in relation to certain aspects of insect sociobiology but not others.

## Conclusion

ORs expanded in the ant ancestor, mainly driven by the 9e subfamily that is associated with social communication. Our analyses detected enrichment of positive selection on ORs in the ancestor of the formicoid ants, mostly in the 9e subfamily, which was accompanied by a further expansion of OR gene numbers. These results support a central role for 9e ORs in the evolution of this large and diverse ant clade. Our study demonstrates the potential of molecular evolutionary analyses to provide insights into the role of ORs in the social evolution and diversification of ants, especially regarding specific sociobiological traits in specific lineages. Detailed phylogenetic dissection of trait evolution reveal a mixture of enrichment of positive selection associated with continuous worker polymorphism and depletion associated with other sociobiological traits. It would be interesting to investigate the underlying causes for the depletion in positive selection in more recent branches, and what explains the differences observed between discrete and continuous worker polymorphism, since the display of discrete worker castes si commonly considered among the most elaborate features of insect sociobiology. Future studies including more species as well as functional genomic data could shed more light on the evolution of these sociobiological traits.

## Supporting information

Supplementary Table S1 - Genomes_ORs data

Supplementary Table S2 - PGLS

Supplementary Table S3 - Positive selection

## Acknowledgements

We thank Sean McKenzie for help with using the *HAPpy-ABCENTH* pipeline for OR annotations. This study was funded by Israel Science Foundation grant number 1845/22.

## Data availability

All the data generated in this study was submitted to the zenodo repository (10.5281/zenodo.16753530) including the OR annotations in GFF and FASTA format, multiple sequence alignments, gene trees, and generax results.

