## Supplementary figures for "Adaptive evolution of odorant receptors is associated with elaborations of social organization in ants"

Figures S1 to S4: Ancestral trait reconstructions made with the R package ape. For figures S2 to S4, circles show the probability for each trait to be the ancestral state.

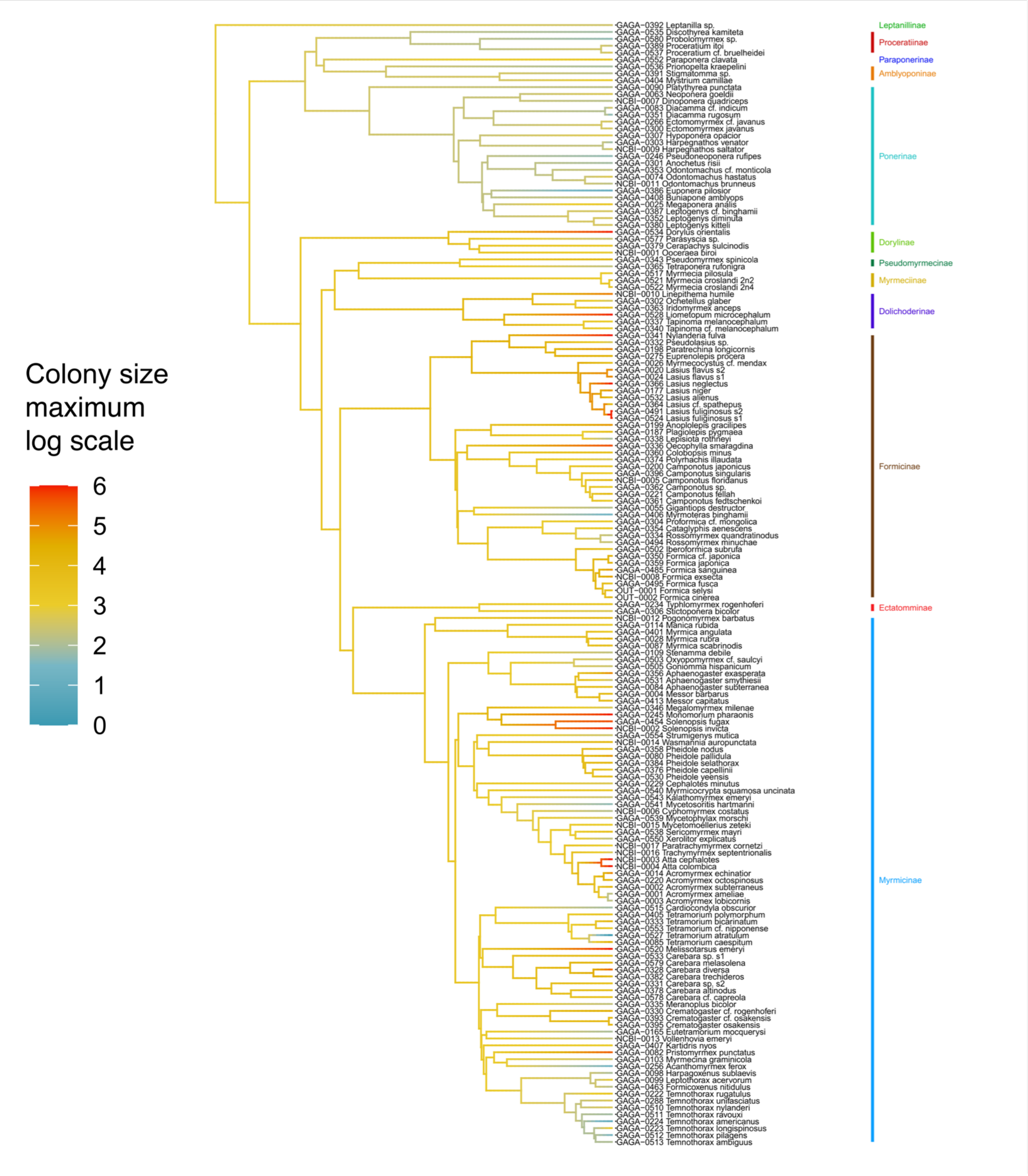


**Figure S1: Ancestral state reconstruction of the log10 maximum colony size.** Ancestral states were inferred using the R package *ape*.

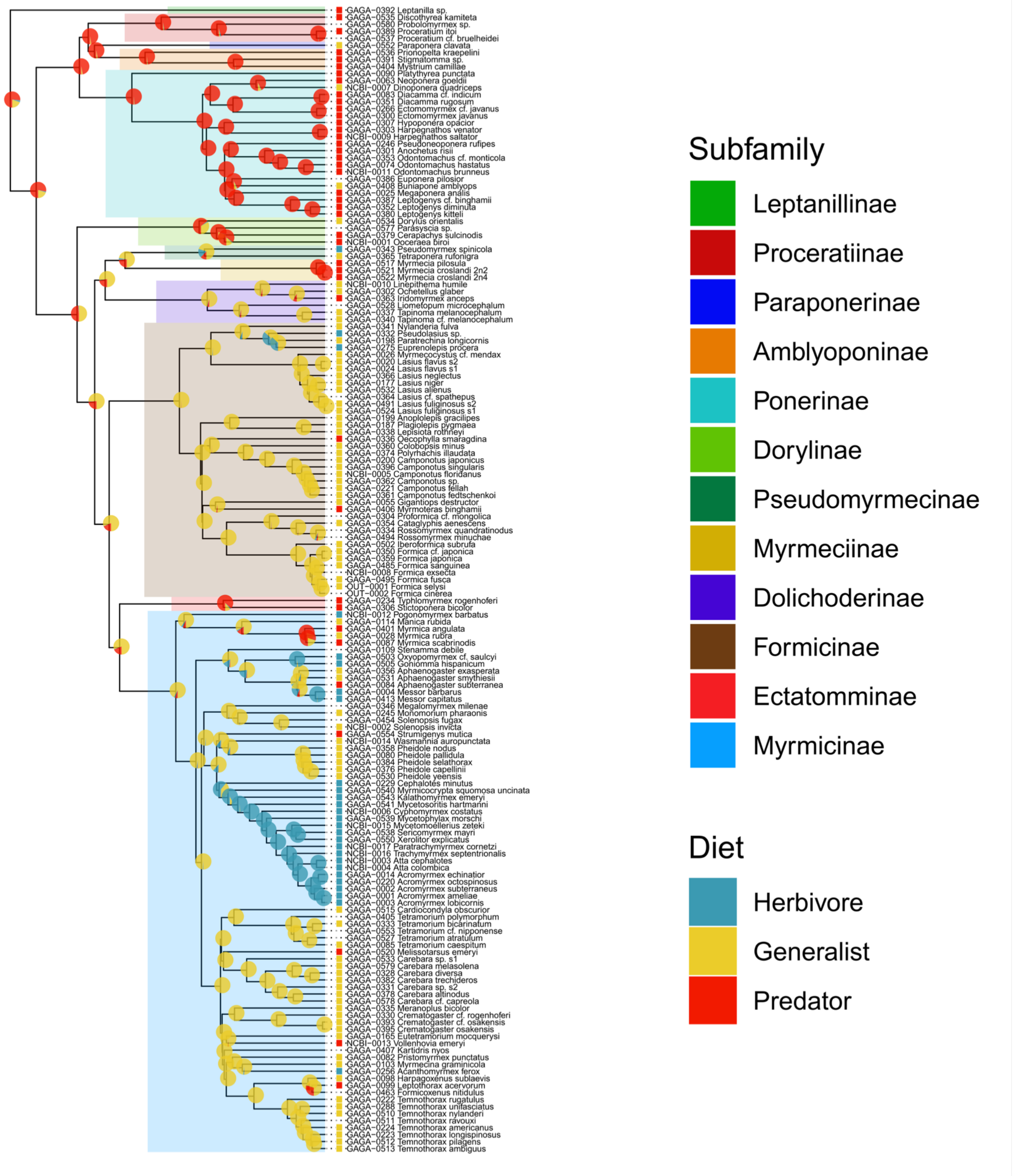


**Figure S2: Ancestral state reconstruction of the diet.** Ancestral states were inferred using the R package *ape*.

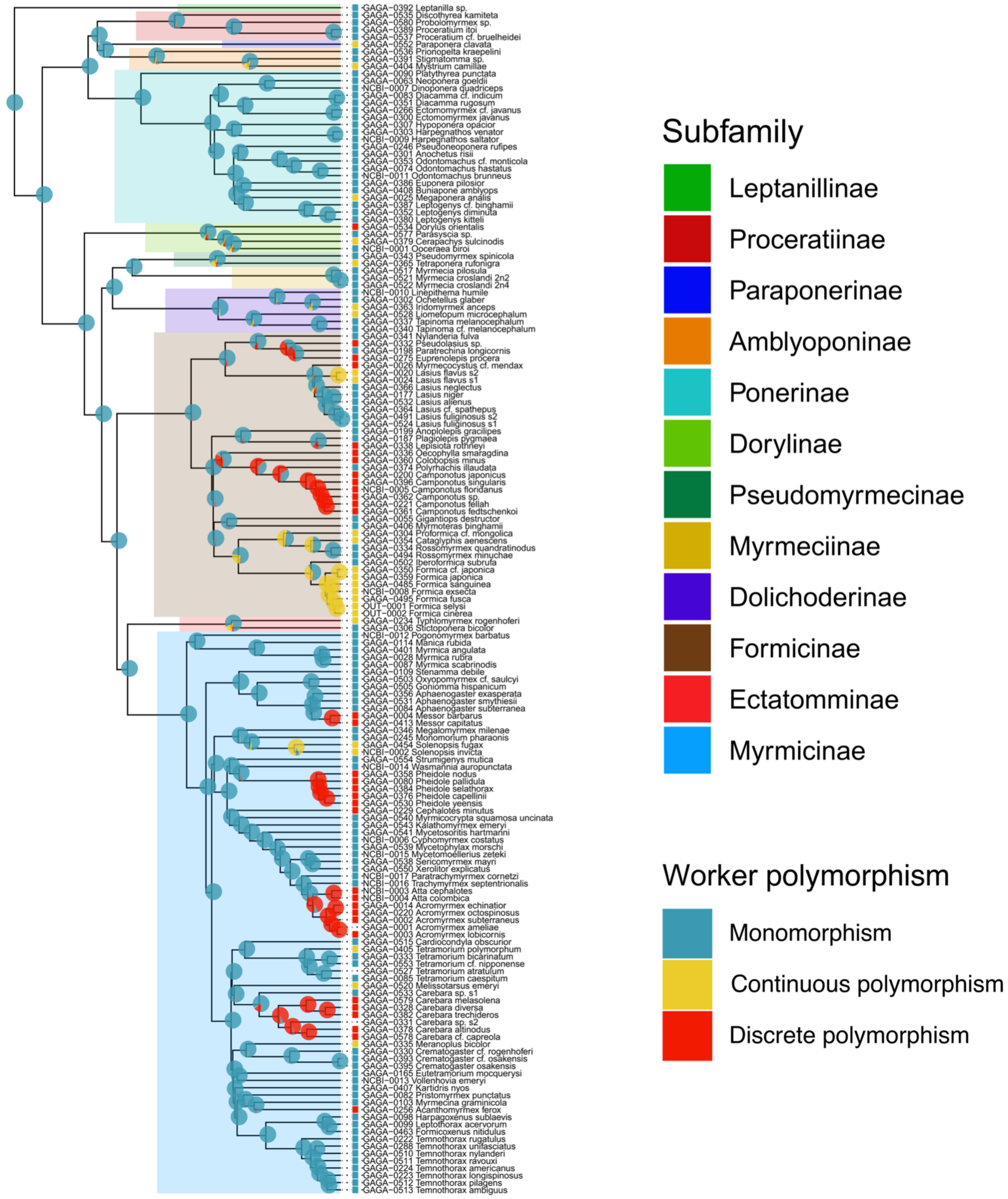


**Figure S3: Ancestral state reconstruction of the worker polymorphism.** Ancestral states were inferred using the R package *ape*.

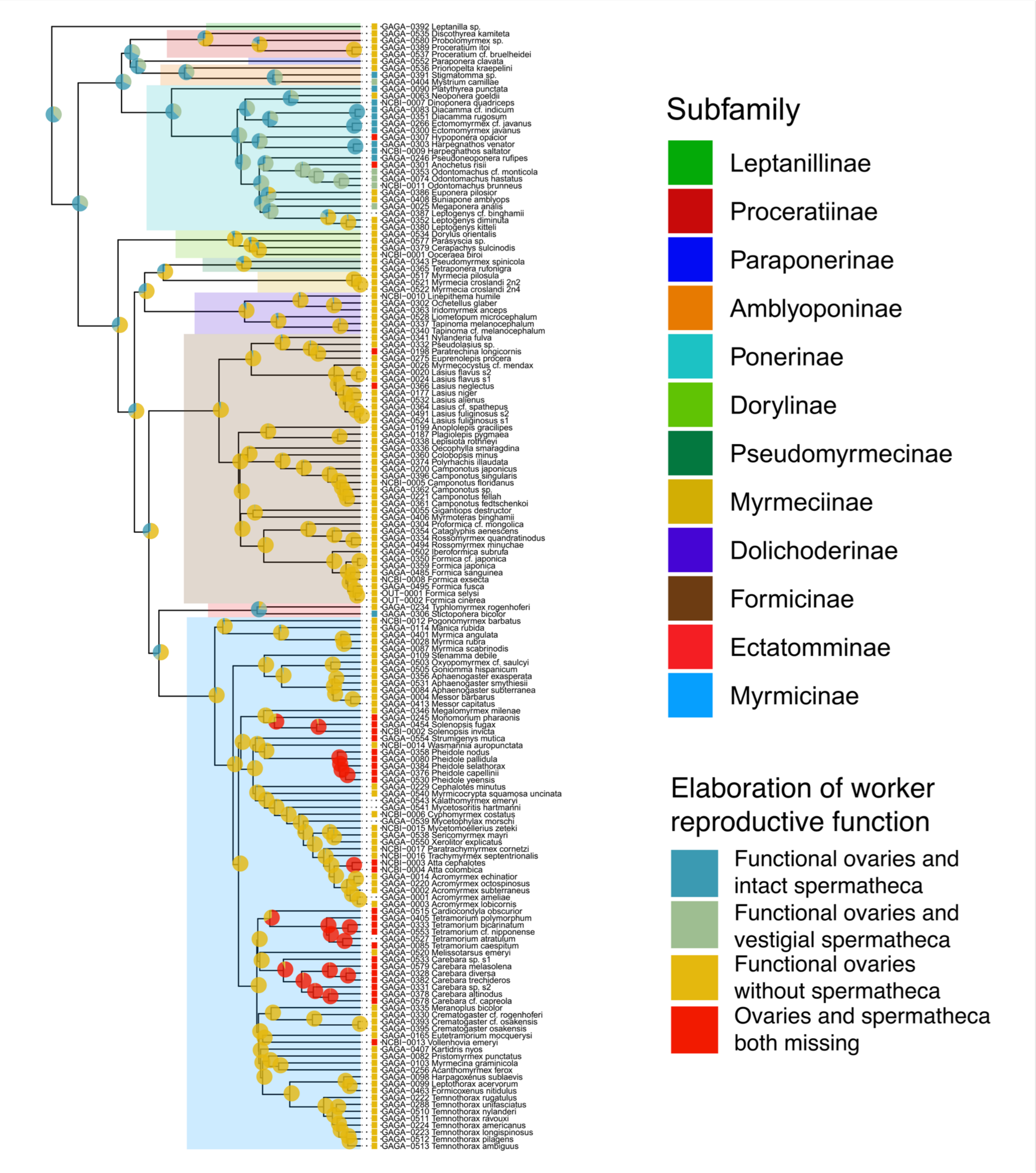


**Figure S4: Ancestral state reconstruction of the worker reproductive function.** Ancestral states were inferred using the R package *ape*.

### Read Segmentation

| Value | Analysis Metric |
| --- | --- |
| 10,878,248 | Reads |
| 86,346,101 | Segmented reads (S-reads) |
| 1,500 | Mean length of S-reads |
| 97.49 % | Percent of reads with full arrays |
| 7.94 | Mean array size (concatenation factor) |

### S-read Length Distribution

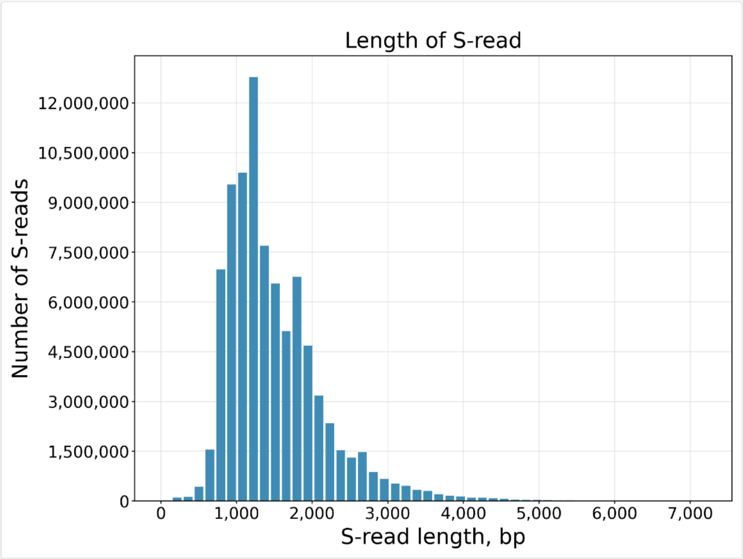


**Figure S5: Overview of the Iso-Seq sequencing results.**

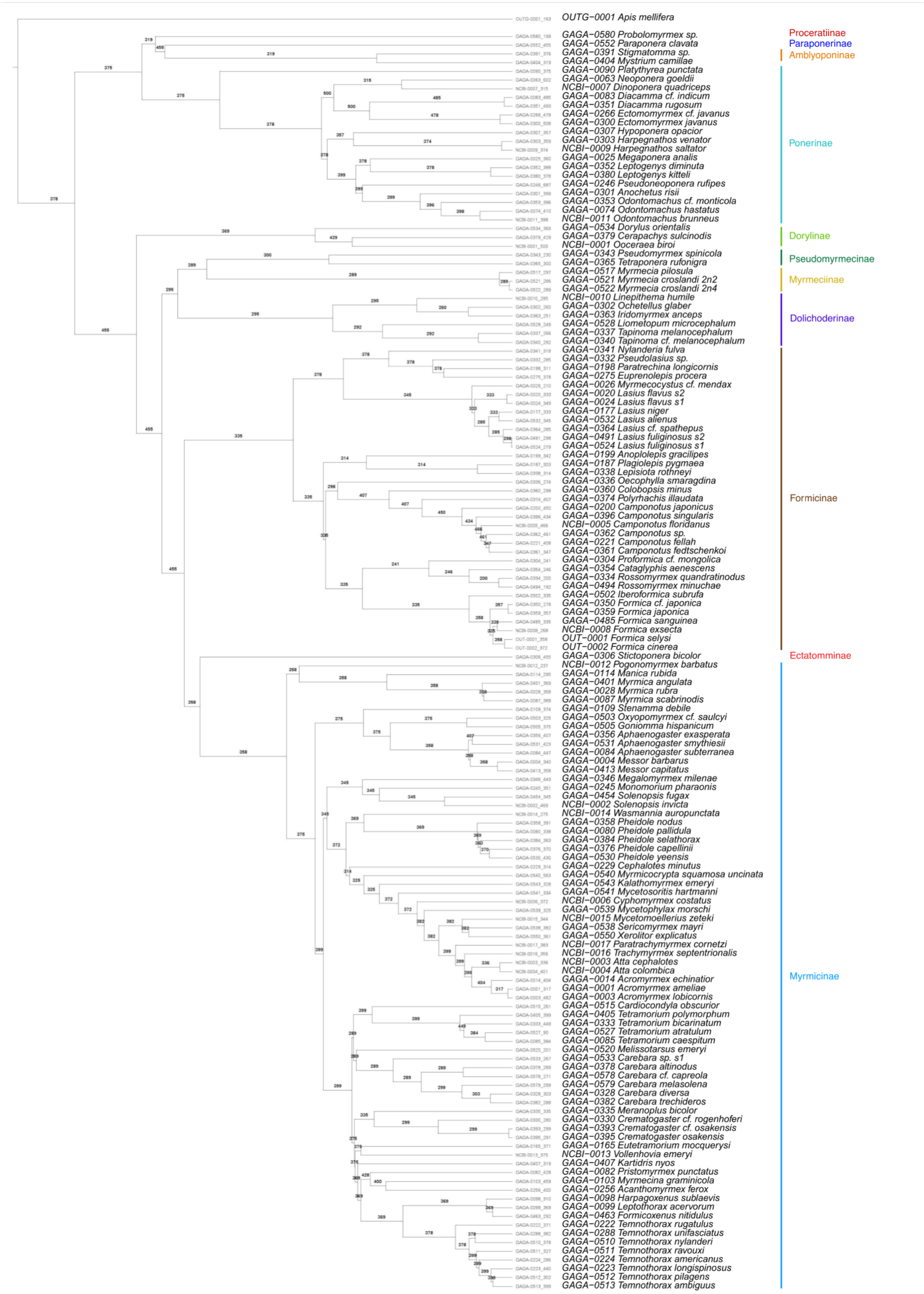


**Figure S6: Inferred OR numbers plotted the ant phylogeny.** The honey bee *Apis mellifera* is used as outgroup for all the ants.

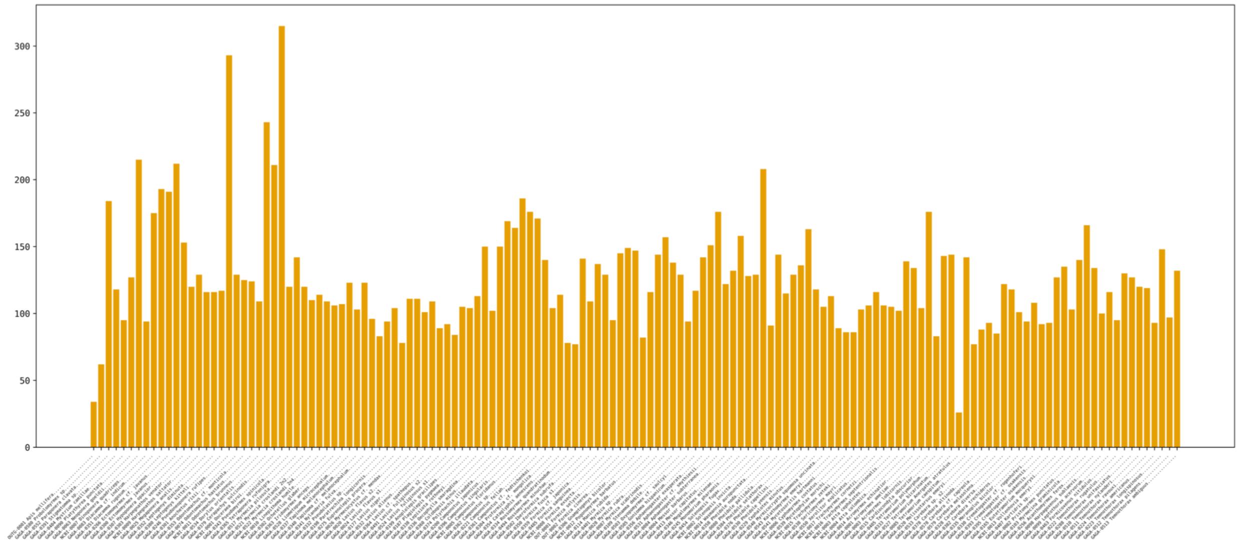


**Figure S7: Number of subfamily 9-exon ORs per species**

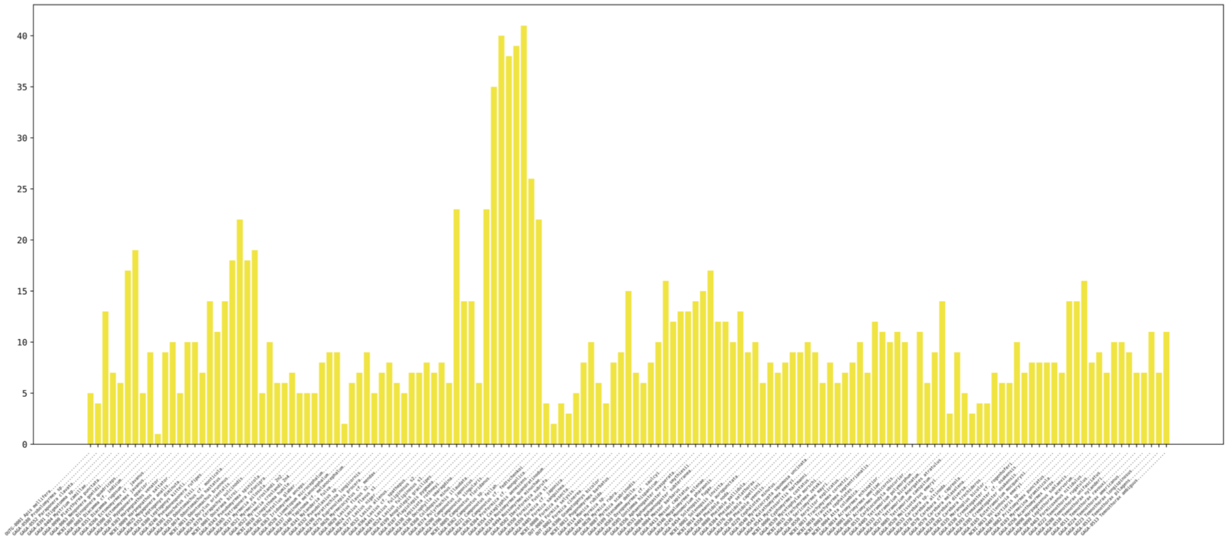


**Figure S8: Number of subfamily P ORs per species**

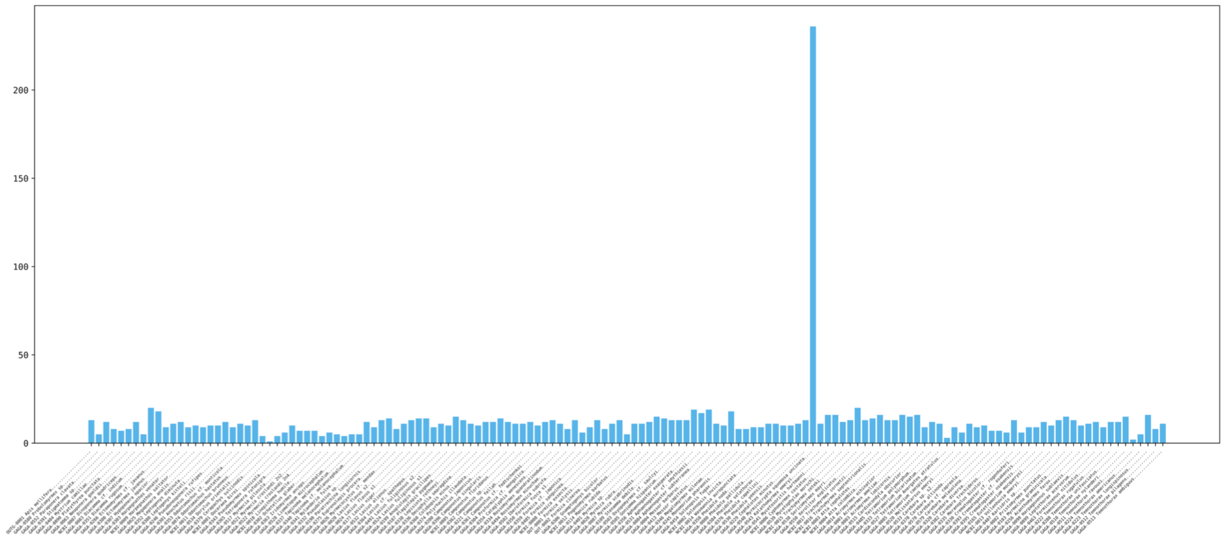


**Figure S9: Number of subfamily H ORs per species**

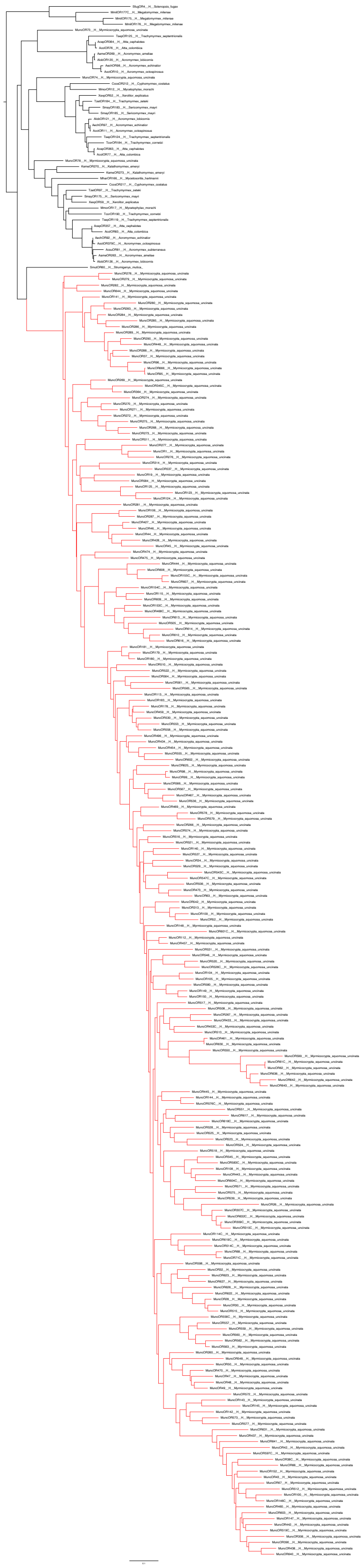

**Figure S10: Phylogenetic tree showing the *Myrmicocrypta squomosa uncinata* specific expansion in the H ORs subfamily in red.**
