## Supplementary Table S2 - PGLS for "Adaptive evolution of odorant receptors is associated with elaborations of social organization in ants"

Table S2: Results of the PGLS analysis, after correction for multiple testing.

Red shows q-values of less than 5%, orange shows q-values of less than 10%.

| ORs subfamily | Average colony size | Social parasitism | Inquilinism | Workers polymorphism | Diet | Workers ovary | Workers spermatheca | Workers sterility |
|---|---|---|---|---|---|---|---|---|
| Total ORs | 0.14512209 | 0.00001490 | 0.00000003 | 0.87830646 | 0.85170400 | 0.93534900 | 1.00000000 | 0.99744600 |
| OR_9.exon | 0.01190480 | 0.00002670 | 0.00000003 | 0.87830646 | 0.30158100 | 1.00000000 | 1.00000000 | 0.99744600 |
| OR_A | 0.72294106 | 0.00304436 | 0.01972397 | 0.85814551 | 0.30189000 | 0.91336400 | 1.00000000 | 0.89163100 |
| OR_D | 0.32007683 | 0.22164471 | 0.27175448 | 0.74541856 | 0.95683800 | 0.14780100 | 1.00000000 | 0.99744600 |
| OR_E | 0.82200192 | 0.05677850 | 0.01803762 | 0.74541856 | 0.86333900 | 0.98625400 | 0.72339500 | 0.89163100 |
| OR_F | 0.03187228 | 0.00188470 | 0.00055200 | 0.74541856 | 0.86333900 | 0.98625400 | 1.00000000 | 0.99744600 |
| OR_G | 0.56884505 | 0.20852422 | 1.00000000 | 0.80409492 | 0.98814400 | 1.00000000 | 1.00000000 | 0.99744600 |
| OR_H | 0.99556976 | 0.47499578 | 0.90865603 | 0.74541856 | 0.30158100 | 0.93534900 | 1.00000000 | 0.99744600 |
| OR_I | 0.70400129 | 0.46300096 | 1.00000000 | 0.74541856 | 0.59035000 | 1.00000000 | 0.90025800 | 0.99744600 |
| OR_J | 0.86109280 | 0.30388827 | 0.25695468 | 0.74541856 | 1.00000000 | 0.98625400 | 1.00000000 | 0.99744600 |
| OR_L | 0.20188714 | 0.00002670 | 0.00000000 | 0.85100253 | 0.90443700 | 0.93534900 | 1.00000000 | 0.99744600 |
| OR_M | 0.84273361 | 0.00002670 | 0.00000006 | 0.74541856 | 0.85343200 | 0.93534900 | 0.90628000 | 0.99744600 |
| OR_N | 0.38632040 | 0.07018520 | 0.00178027 | 0.98077757 | 0.54826000 | 0.91336400 | 1.00000000 | 0.99744600 |
| OR_O | 0.02534427 | 0.05677850 | 0.00000101 | 0.74541856 | 0.85343200 | 0.93534900 | 1.00000000 | 0.99744600 |
| OR_P | 0.29849008 | 0.00606160 | 0.00088900 | 0.93526994 | 0.91324400 | 0.98625400 | 1.00000000 | 0.99744600 |
| OR_Q | 0.95748658 | 0.72642294 | 1.00000000 | 0.74541856 | 0.85343200 | 0.93534900 | 1.00000000 | 0.99744600 |
| OR_T | 0.00872355 | 0.01103333 | 0.00023900 | 0.98555930 | 0.59035000 | 0.93534900 | 1.00000000 | 0.99744600 |
| OR_U | 0.56884505 | 0.07828524 | 0.05202740 | 0.91887268 | 0.61700400 | 0.98625400 | 0.84243100 | 0.89163100 |
| OR_V | 0.01190480 | 0.00002670 | 0.00023900 | 0.74541856 | 0.85170400 | 0.98625400 | 1.00000000 | 0.99744600 |
| OR_with_partial | 0.00169118 | 0.00002670 | 0.00000000 | 0.95007586 | 0.86333900 | 0.98625400 | 1.00000000 | 0.99744600 |
