## Supplementary Table S3 - Positive selection for "Adaptive evolution of odorant receptors is associated with elaborations of social organization in ants"

### All ORs

| Clade | Species branch of interest | Sum of the length of all gene tree branches in the clade | Total number of positive gene tree branches in the clade | Sum of the length of gene tree branches mapped on the species branch of interest | Number of positive gene tree branches mapped on the species branch of interest | p-value | q-value | Fold change of normalized positive branches | Trait change mapped on the species branch |
|---|---|---|---|---|---|---|---|---|---|
| **Lasius-Nylanderia** | 68 | 5667 | 276 | 370 | 16 | 0.091620296 | 0.11689486 | 0.887896592 | Increased colony size |
| **Lasius-Nylanderia** | 69 | | | 423 | 18 | 0.081515848 | 0.11689486 | 0.87372803 | Worker Polymorphism (Discrete) |
| **Lasius-Nylanderia** | 92 | | | 207 | 15 | 0.034553758 | 0.11689486 | 1.487870195 | Worker Polymorphism (Continuous) |
| **Lasius-Nylanderia** | 94 | | | 309 | 13 | 0.097889054 | 0.117886164 | 0.863831434 | Diet - Omnivore to Herbivore, Worker Polymorphism (Discrete) |
| **Lasius-Nylanderia** | 110 | | | 113 | 14 | 0.000735556 | 0.024611531 | 2.543863024 | Worker Polymorphism (Continuous) |
| **Lasius-Nylanderia** | 112 | | | 305 | 10 | 0.048421076 | 0.11689486 | 0.673200285 | Diet - Omnivore to Herbivore, Worker Polymorphism (Discrete) |
| **Lasius-Nylanderia** | 179 | | | 36 | 2 | 0.274611028 | 0.274611028 | 1.140700483 | Increased colony size |
| **Leaf-cutter-ants** | 305 | 9260 | 344 | 372 | 9 | 0.048342252 | 0.11689486 | 0.651256564 | Worker Polymorphism (Discrete) |
| **Leaf-cutter-ants** | 311 | | | 319 | 8 | 0.067298973 | 0.11689486 | 0.675074725 | Increased colony size, Worker Polymorphism (Discrete) |
| **Leaf-cutter-ants** | 314 | | | 294 | 8 | 0.090454767 | 0.11689486 | 0.732479038 | Increased colony size, Loss of Worker Reproductive Function, Worker Polymorphism (Discrete) |
| **Leaf-cutter-ants** | 315 | | | 325 | 8 | 0.062176311 | 0.11689486 | 0.662611807 | Increased colony size, Worker Polymorphism (Discrete) |
| **Poneroid** | 12 | 4827 | 271 | 274 | 8 | 0.013470676 | 0.098042717 | 0.520052792 | Loss of Worker Reproductive Function |
| **Formica-Camponotus** | 97 | 13301 | 495 | 563 | 10 | 0.002865778 | 0.032618275 | 0.477277213 | Worker Polymorphism (Discrete) |
| **Formica-Camponotus** | 115 | | | 353 | 13 | 0.113434837 | 0.123443793 | 0.989572782 | Worker Polymorphism (Continuous) |
| **Formica-Camponotus** | 116 | | | 642 | 22 | 0.081118866 | 0.11689486 | 0.920803046 | Worker Polymorphism (Discrete) |
| **Formica-Camponotus** | 138 | | | 285 | 6 | 0.046836174 | 0.11689486 | 0.565699096 | Worker Polymorphism (Continuous) |
| **Formica-Camponotus** | 139 | | | 358 | 17 | 0.060849178 | 0.11689486 | 1.275983297 | Increased colony size, Worker Polymorphism (Continuous) |
| **Formica-Camponotus** | 140 | | | 444 | 11 | 0.040045334 | 0.11689486 | 0.665715716 | Worker Polymorphism (Discrete) |
| **Formica-Camponotus** | 157 | | | 184 | 5 | 0.133999282 | 0.137721484 | 0.730182257 | Worker Polymorphism (Continuous) |
| **Formica-Camponotus** | 158 | | | 195 | 10 | 0.079378381 | 0.11689486 | 1.377984978 | Increased colony size, Worker Polymorphism (Continuous) |
| **Formica-Camponotus** | 160 | | | 440 | 20 | 0.061933249 | 0.11689486 | 1.221395776 | Worker Polymorphism (Discrete) |
| **Messor** | 144 | 6227 | 265 | 466 | 23 | 0.06765569 | 0.11689486 | 1.15977812 | Increased colony size |
| **Messor** | 145 | | | 430 | 18 | 0.098769489 | 0.117886164 | 0.983641948 | Diet - Omnivore to Herbivore |
| **Messor** | 186 | | | 325 | 11 | 0.088208717 | 0.11689486 | 0.795320755 | Diet - Omnivore to Herbivore |
| **Pheidole** | 172 | 9350 | 467 | 442 | 21 | 0.088283424 | 0.11689486 | 0.951243617 | Loss of Worker Reproductive Function |
| **Pheidole** | 199 | | | 470 | 32 | 0.015898819 | 0.098042717 | 1.363160053 | Worker Polymorphism (Continuous) |
| **Pheidole** | 200 | | | 487 | 13 | 0.0035263 | 0.032618275 | 0.53445251 | Increased colony size |
| **Pheidole** | 228 | | | 512 | 33 | 0.024465661 | 0.11689486 | 1.290442653 | Increased colony size, Loss of Worker Reproductive Function, Worker Polymorphism (Discrete) |
| **Carebara** | 171 | 8531 | 460 | 716 | 23 | 0.001330353 | 0.024611531 | 0.595740223 | Loss of Worker Reproductive Function |
| **Carebara** | 194 | | | 284 | 16 | 0.102242077 | 0.118217402 | 1.044825475 | Loss of Worker Reproductive Function, Worker Polymorphism (Discrete) |
| **Carebara** | 222 | | | 190 | 11 | 0.120870277 | 0.12777715 | 1.073695652 | Worker Polymorphism (Discrete) |
| **Myrmica** | 120 | 3731 | 245 | 288 | 24 | 0.042528426 | 0.11689486 | 1.269047619 | Diet - Omnivorevore to Carnivore |
| **Leaf-cutter-ancestor** | 175 | 11946 | 537 | 509 | 16 | 0.029218855 | 0.11689486 | 0.699278902 | Diet - Omnivore to Herbivore |
| **Leaf-cutter-ancestor** | 203 | | | 337 | 12 | 0.080724889 | 0.11689486 | 0.79213567 | Diet - Omnivore to Herbivore |
| **Formicoid** | 5 | 16953 | 981 | 337 | 21 | 0.085158892 | 0.11689486 | 1.076879101 | |
| **Formicoid** | 9 | | | 327 | 22 | 0.068521074 | 0.11689486 | 1.162659335 | |
| **Formicoid** | 17 | | | 252 | 14 | 0.108464326 | 0.121611517 | 0.960074754 | |

### 9-Exons only

| Clade | Species branch | Sum of the length of all gene tree branches | Total number of positive gene tree branches | Sum of the length of gene tree branches mapped on the species branch of interest | Number of positive gene tree branches mapped on the species branch of interest | p-value | q-value | Fold change of normalized positive branches | Trait change mapped on the species branch |
|---|---|---|---|---|---|---|---|---|---|
| **Lasius-Nylanderia** | 68 | | | 117 | 8 | 0.148437 | 0.192875714 | 0.962691629 | Increased colony size |
| **Lasius-Nylanderia** | 69 | | | 126 | 6 | 0.090181 | 0.187753857 | 0.670445956 | Worker Polymorphism (Discrete) |
| **Lasius-Nylanderia** | 92 | | | 58 | 9 | 0.012342 | 0.152218 | 2.184729064 | Worker Polymorphism (Continuous) |
| **Lasius-Nylanderia** | 94 | 1774 | 126 | 96 | 6 | 0.160439 | 0.192875714 | 0.879960317 | Diet - Omnivore to Herbivore, Worker Polymorphism (Discrete) |
| **Lasius-Nylanderia** | 110 | | | 40 | 10 | 0.000251 | 0.009287 | 3.51984127 | Worker Polymorphism (Continuous) |
| **Lasius-Nylanderia** | 112 | | | 100 | 6 | 0.15328 | 0.192875714 | 0.844761905 | Diet - Omnivore to Herbivore, Worker Polymorphism (Discrete) |
| **Lasius-Nylanderia** | 179 | | | 14 | 0 | 0.35509 | 0.35509 | #NUM! | Increased colony size |
| **Leaf-cutter-ants** | 305 | | | 138 | 4 | 0.099643 | 0.187753857 | 0.594902549 | Worker Polymorphism (Discrete) |
| **Leaf-cutter-ants** | 311 | 2976 | 145 | 116 | 3 | 0.102116 | 0.187753857 | 0.530796671 | Increased colony size, Worker Polymorphism (Discrete) |
| **Leaf-cutter-ants** | 314 | | | 97 | 5 | 0.181793 | 0.192875714 | 1.057945254 | Increased colony size, Loss of Worker Reproductive Function, Worker Polymorphism (Discrete) |
| **Leaf-cutter-ants** | 315 | | | 110 | 5 | 0.180337 | 0.192875714 | 0.932915361 | Increased colony size, Worker Polymorphism (Discrete) |
| **Poneroid** | 12 | 1547 | 114 | 72 | 4 | 0.168141 | 0.192875714 | 0.753898635 | Loss of Worker Reproductive Function |
| **Formica-Camponotus** | 97 | | | 207 | 6 | 0.076449 | 0.187753857 | 0.636832212 | Worker Polymorphism (Discrete) |
| **Formica-Camponotus** | 115 | | | 145 | 7 | 0.153032 | 0.192875714 | 1.060655028 | Worker Polymorphism (Continuous) |
| **Formica-Camponotus** | 116 | | | 231 | 11 | 0.12372 | 0.192875714 | 1.046224347 | Worker Polymorphism (Discrete) |
| **Formica-Camponotus** | 138 | | | 130 | 2 | 0.043386 | 0.187753857 | 0.338010943 | Worker Polymorphism (Continuous) |
| **Formica-Camponotus** | 139 | 5251 | 239 | 148 | 9 | 0.09536 | 0.187753857 | 1.336056768 | Increased colony size, Worker Polymorphism (Continuous) |
| **Formica-Camponotus** | 140 | | | 201 | 8 | 0.133534 | 0.192875714 | 0.874456171 | Worker Polymorphism (Discrete) |
| **Formica-Camponotus** | 157 | | | 98 | 5 | 0.176003 | 0.192875714 | 1.120954658 | Worker Polymorphism (Continuous) |
| **Formica-Camponotus** | 158 | | | 87 | 7 | 0.056573 | 0.187753857 | 1.76775838 | Increased colony size, Worker Polymorphism (Continuous) |
| **Formica-Camponotus** | 160 | | | 189 | 11 | 0.089983 | 0.187753857 | 1.278718647 | Worker Polymorphism (Discrete) |
| **Messor** | 144 | | | 161 | 8 | 0.14779 | 0.192875714 | 1.041603188 | Increased colony size |
| **Messor** | 145 | 2222 | 106 | 146 | 9 | 0.10432 | 0.187753857 | 1.292194365 | Diet - Omnivore to Herbivore |
| **Messor** | 186 | | | 122 | 7 | 0.140003 | 0.192875714 | 1.202752861 | Diet - Omnivore to Herbivore |
| **Pheidole** | 172 | | | 134 | 6 | 0.145517 | 0.192875714 | 0.820008068 | Loss of Worker Reproductive Function |
| **Pheidole** | 199 | 3388 | 185 | 184 | 15 | 0.033401 | 0.187753857 | 1.492949471 | Worker Polymorphism (Continuous) |
| **Pheidole** | 200 | | | 179 | 4 | 0.018186 | 0.1682205 | 0.409240525 | Increased colony size |
| **Pheidole** | 228 | | | 190 | 12 | 0.106563 | 0.187753857 | 1.156642959 | Increased colony size, Loss of Worker Reproductive Function, Worker Polymorphism (Discrete) |
| **Carebara** | 171 | | | 249 | 5 | 0.003247 | 0.0600695 | 0.367359007 | Loss of Worker Reproductive Function |
| **Carebara** | 194 | 2982 | 163 | 79 | 4 | 0.200383 | 0.205949194 | 0.92630271 | Loss of Worker Reproductive Function, Worker Polymorphism (Discrete) |
| **Carebara** | 222 | | | 65 | 1 | 0.095583 | 0.187753857 | 0.281453516 | Worker Polymorphism (Discrete) |
| **Myrmica** | 120 | 1461 | 102 | 117 | 12 | 0.050246 | 0.187753857 | 1.46907994 | Diet - Omnivorevore to Carnivore |
| **Leaf-cutter-ancestor** | 175 | 3762 | 196 | 157 | 5 | 0.082799 | 0.187753857 | 0.611269986 | Diet - Omnivore to Herbivore |
| **Leaf-cutter-ancestor** | 203 | | | 97 | 5 | 0.18245 | 0.192875714 | 0.989375131 | Diet - Omnivore to Herbivore |
| **Formicoid** | 5 | | | 112 | 9 | 0.098298 | 0.187753857 | 1.320283883 | |
| **Formicoid** | 9 | 5767 | 351 | 108 | 10 | 0.057171 | 0.187753857 | 1.521314762 | |
| **Formicoid** | 17 | | | 79 | 7 | 0.097779 | 0.187753857 | 1.455840456 | |

### Without 9-Exons

| Clade | Species branch | Sum of the length of all gene tree branches | Total number of positive gene tree branches | Sum of the length of gene tree branches mapped on the species branch of interest | Number of positive gene tree branches mapped on the species branch of interest | p-value | q-value | Fold change of normalized positive branches | Trait change mapped on the species branch |
|---|---|---|---|---|---|---|---|---|---|
| **Lasius-Nylanderia** | 68 | 3893 | 150 | 253 | 8 | 0.121234 | 0.161843103 | 0.820658762 | Increased colony size |
| **Lasius-Nylanderia** | 69 | | | 299 | 12 | 0.120345 | 0.161843103 | 1.041605351 | Worker Polymorphism (Discrete) |
| **Lasius-Nylanderia** | 92 | | | 150 | 6 | 0.166553 | 0.176070314 | 1.038133333 | Worker Polymorphism (Continuous) |
| **Lasius-Nylanderia** | 94 | | | 214 | 7 | 0.139209 | 0.1716911 | 0.84894081 | Diet - Omnivore to Herbivore, Worker Polymorphism (Discrete) |
| **Lasius-Nylanderia** | 110 | | | 72 | 4 | 0.157994 | 0.174522471 | 1.441851852 | Worker Polymorphism (Continuous) |
| **Lasius-Nylanderia** | 112 | | | 205 | 4 | 0.056159 | 0.161843103 | 0.506406504 | Diet - Omnivore to Herbivore, Worker Polymorphism (Discrete) |
| **Lasius-Nylanderia** | 179 | | | 21 | 2 | 0.148012 | 0.174522471 | 2.471746032 | Increased colony size |
| **Leaf-cutter-ants** | 305 | 6284 | 199 | 227 | 5 | 0.120841 | 0.161843103 | 0.695548226 | Worker Polymorphism (Discrete) |
| **Leaf-cutter-ants** | 311 | | | 199 | 5 | 0.154353 | 0.174522471 | 0.793414308 | Increased colony size, Worker Polymorphism (Discrete) |
| **Leaf-cutter-ants** | 314 | | | 197 | 3 | 0.07594 | 0.161843103 | 0.480881565 | Increased colony size, Loss of Worker Reproductive Function, Worker Polymorphism (Discrete) |
| **Leaf-cutter-ants** | 315 | | | 213 | 3 | 0.056885 | 0.161843103 | 0.444759006 | Increased colony size, Worker Polymorphism (Discrete) |
| **Poneroid** | 12 | 3280 | 157 | 204 | 4 | 0.018213 | 0.161843103 | 0.409641564 | Loss of Worker Reproductive Function |
| **Formica-Camponotus** | 97 | 8050 | 256 | 359 | 4 | 0.006512 | 0.161843103 | 0.350365599 | Worker Polymorphism (Discrete) |
| **Formica-Camponotus** | 115 | | | 207 | 6 | 0.160372 | 0.174522471 | 0.911458333 | Worker Polymorphism (Continuous) |
| **Formica-Camponotus** | 116 | | | 414 | 11 | 0.100814 | 0.161843103 | 0.835503472 | Worker Polymorphism (Discrete) |
| **Formica-Camponotus** | 138 | | | 153 | 4 | 0.183086 | 0.188171722 | 0.822099673 | Worker Polymorphism (Continuous) |
| **Formica-Camponotus** | 139 | | | 210 | 8 | 0.12685 | 0.161843103 | 1.197916667 | Increased colony size, Worker Polymorphism (Continuous) |
| **Formica-Camponotus** | 140 | | | 240 | 3 | 0.033219 | 0.161843103 | 0.393066406 | Worker Polymorphism (Discrete) |
| **Formica-Camponotus** | 157 | | | 82 | 0 | 0.069689 | 0.161843103 | #NUM! | Worker Polymorphism (Continuous) |
| **Formica-Camponotus** | 158 | | | 107 | 3 | 0.222739 | 0.222739 | 0.881644276 | Increased colony size, Worker Polymorphism (Continuous) |
| **Formica-Camponotus** | 160 | | | 249 | 9 | 0.12667 | 0.161843103 | 1.13657756 | Worker Polymorphism (Discrete) |
| **Messor** | 144 | 4005 | 159 | 306 | 15 | 0.077427 | 0.161843103 | 1.234739179 | Increased colony size |
| **Messor** | 145 | | | 285 | 9 | 0.103864 | 0.161843103 | 0.795431976 | Diet - Omnivore to Herbivore |
| **Messor** | 186 | | | 203 | 4 | 0.051556 | 0.161843103 | 0.496328655 | Diet - Omnivore to Herbivore |
| **Pheidole** | 172 | 5962 | 282 | 306 | 15 | 0.10682 | 0.161843103 | 1.036364901 | Loss of Worker Reproductive Function |
| **Pheidole** | 199 | | | 287 | 17 | 0.065646 | 0.161843103 | 1.252304347 | Worker Polymorphism (Continuous) |
| **Pheidole** | 200 | | | 308 | 9 | 0.03537 | 0.161843103 | 0.617781155 | Increased colony size |
| **Pheidole** | 228 | | | 323 | 21 | 0.031739 | 0.161843103 | 1.374547131 | Increased colony size, Loss of Worker Reproductive Function, Worker Polymorphism (Discrete) |
| **Carebara** | 171 | 5549 | 297 | 467 | 18 | 0.028664 | 0.161843103 | 0.720134969 | Loss of Worker Reproductive Function |
| **Carebara** | 194 | | | 208 | 12 | 0.115978 | 0.161843103 | 1.077894328 | Loss of Worker Reproductive Function, Worker Polymorphism (Discrete) |
| **Carebara** | 222 | | | 125 | 10 | 0.06103 | 0.161843103 | 1.494680135 | Worker Polymorphism (Discrete) |
| **Myrmica** | 120 | 2270 | 143 | 171 | 12 | 0.114544 | 0.161843103 | 1.113973746 | Diet - Omnivorevore to Carnivore |
| **Leaf-cutter-ancestor** | 175 | 8184 | 341 | 353 | 11 | 0.070807 | 0.161843103 | 0.747875354 | Diet - Omnivore to Herbivore |
| **Leaf-cutter-ancestor** | 203 | | | 241 | 7 | 0.088141 | 0.161843103 | 0.697095436 | Diet - Omnivore to Herbivore |
| **Formicoid** | 5 | 11186 | 630 | 225 | 12 | 0.116379 | 0.161843103 | 0.946962963 | |
| **Formicoid** | 9 | | | 219 | 12 | 0.118187 | 0.161843103 | 0.972907154 | |
| **Formicoid** | 17 | | | 173 | 7 | 0.096769 | 0.161843103 | 0.718432884 | |
